# Agricultural land use induces broader homogenization of soil microbial functional composition than taxonomic composition

**DOI:** 10.1101/2025.02.17.638669

**Authors:** Takamitsu Ohigashi, Yvonne M. Madegwa, George N. Karuku, Keston Njira, Yoshitaka Uchida

## Abstract

Land-use changes from natural ecosystems to farmlands significantly alter soil functioning worldwide, especially challenging sub-Saharan Africa with rapid population growth and intensive agriculture. Soil microbial diversity is vital in supporting ecosystem multifunctionality and preventing pathogen growth. Recent studies have revealed that farming activities homogenize microbial communities across distant sites, which may lead to functional homogenization on that scale. However, given the redundancy of microbial functions, functional homogenization driven by farming may occur on a broader scale than taxonomic homogenization. We compared the taxonomic and functional compositions of soil prokaryotic and fungal communities between natural lands and farmlands at scales ranging from within-site (∼200 m) to across-site (∼1500 km) in Kenya and Malawi, using amplicon sequencing of 16S rRNA and ITS genes and the prediction of microbial functions. Soil microbial functional compositions were homogenized more broadly than taxonomic compositions in farmlands compared to natural lands, suggesting that similar functional responses to farming occur across scales where different taxa thrive. Furthermore, environmental factors predominantly influenced within-site homogeneity, whereas farming itself was a significant contributor to across-site homogeneity, indicating an overriding influence of farming compared to environmental variations. Additionally, pathogenic fungi were relatively more abundant in farmlands, likely due to reduced species competition and farming-induced environmental changes such as low soil pH. Our findings highlight the need to investigate microbial functional diversity alongside taxonomic diversity when assessing the impacts of land-use changes on soil health to develop sustainable land management strategies.

## Introduction

Maintaining soil health during land-use changes from natural ecosystems to farmlands is a global challenge (Kibblewhite *et al*. 2007). Land-use changes and resulting intensive farming practices are causing a serious threat, declining soil nutrients, particularly in sub-Saharan Africa (Sanchez 2002; Tully *et al*. 2015). The depletion of soil nutrients can impact soil microbial diversity, which has been shown to enhance soil multifunctionality, including chemical cycling and pathogen resistance (Elsas *et al*. 2012; Delgado-Baquerizo *et al*. 2016; Romero *et al*. 2023). Therefore, maintaining diverse microbial species while sustaining agricultural productivity is key to sustainable development. However, information on soil microbial communities in this region remains limited, even though soil degradation is severe (Makhalanyane *et al*. 2023). Considering soil ecology dynamics alongside soil chemical and physical properties is essential to addressing the challenges of sustainable land management (Bender *et al*. 2016).

Land-use changes associated with cultivation practices, including weed removal and monoculture, generally decrease soil nutrients such as soil carbon (C) (Tully *et al*. 2015; Ohigashi *et al*. 2021). Yet these practices act as moderate disturbances that suppress the dominance of certain species and create niches for others, leading to an increase in microbial sample-scale diversity (*α* diversity) as observed in various regions such as Europe, South America, and Africa (Tardy *et al*. 2015; Ohigashi *et al*. 2021; Labouyrie *et al*. 2023). Conversely, reduced microbial community heterogeneity (*β* diversity) within sites was observed due to the homogenization of environmental factors (e.g., soil chemical property) by land-use changes, including agriculture and urbanization (Rodrigues *et al*. 2013; Tatsumi *et al*. 2023a). This observation suggests that even though sample-scale microbial diversity may increase by land-use changes, microbial community compositions can become similar among different scoops of soil within the converted land.

Moreover, the influences of land-use changes on microbial communities can extend not only within one site but also across broader geographic scales, potentially spanning a specific region (e.g., sub-Saharan Africa). Generally, ecological communities, including soil microbes, exhibit greater dissimilarity with increasing geographical distance, known as the distance-decay relationship (Hanson *et al*. 2012; Nemergut *et al*. 2013; Socolar *et al*. 2016; Li *et al*. 2020). This pattern is primarily driven by environmental variations, species dispersal limitations, and local historical factors associated with the geographical distance (Hanson *et al*. 2012). However, farming activities, which simultaneously homogenize landscapes at multiple sites, can lead to microbial community homogenization even among distant sites (Banerjee *et al*. 2024; Peng *et al*. 2024). A meta-analysis using over 2,400 soil samples across six continents revealed a global homogenization of prokaryotic communities in croplands compared to forests, grasslands, and wetlands (Peng *et al*. 2024). This taxonomic homogenization may reflect the widespread influence of farming, but its ecological significance lies in how these taxonomic changes are expressed as shifts in microbial functions, which are critical for soil health.

Taxonomic changes in microbial communities are often correlated with their functional changes across multiple sites (Liang *et al*. 2023; Peng *et al*. 2024), suggesting that taxonomic homogenization can lead to functional homogenization. In contrast, some studies have reported mismatches between variations in taxonomic and functional compositions (Langenheder *et al*. 2005, 2006; Knight *et al*. 2024). Laboratory incubations using bacterial communities from different sources revealed similar functional patterns (e.g., biomass production and respiration) despite taxonomic dissimilarity (Langenheder *et al*. 2005, 2006). Similarly, prokaryotic and fungal functional responses to extreme conditions were relatively consistent across soil microcosms collected throughout Europe, while taxonomic responses varied depending on local factors such as native climate (Knight *et al*. 2024). Thus, although microbial taxonomic compositions vary from site to site because of the limited distributions of certain taxa, functional responses to environmental changes may be similar due to functional redundancy among different taxa (Louca *et al*. 2018). As agricultural practices simultaneously homogenize landscapes, they may act as selective pressures on certain soil microbial functions rather than taxa.

Additionally, such an imbalanced microbiome may threaten soil health. For instance, microbial communities with decreased taxonomic and functional diversity tend to allow invader species to increase, including pathogenic microbes (Elsas *et al*. 2012; Eisenhauer *et al*. 2013). However, whether broader-scale microbial community homogeneity (i.e., homogeneity across farmlands) enables the widespread distribution of pathogens remains unclear. Understanding the ecological mechanisms underlying interactions between pathogens and other species is critical for sustainable land management (Johnson *et al*. 2015).

Many studies have explored the factors influencing microbial community similarity or dissimilarity across sites, such as environmental and geographic patterns (Kuramae *et al*. 2012; Tedersoo *et al*. 2014; Delgado-Baquerizo *et al*. 2017, 2018; Zhang *et al*. 2020b). Nevertheless, the underlying mechanisms of community assembly remain under discussion. Community assembly can be driven by four primary ecological processes: environmental selection, dispersal, speciation, and ecological drift (Vellend 2010). The relative importance of these processes depends on the geographic scales, organism body size, and ecological traits (Martiny *et al*. 2011; Farjalla *et al*. 2012; Tecon & Or 2017). For instance, due to their smaller body size, prokaryotic assemblies are more influenced by stochastic processes than fungal assemblies (Luan *et al*. 2020). Meanwhile, some fungi form spores that allow for relatively efficient dispersal, even across continents (Tedersoo *et al*. 2014). Moreover, strong environmental stressors, such as extreme soil pH or drought, can amplify the role of environmental selection (Tripathi *et al*. 2018; Bei *et al*. 2023; Ning *et al*. 2024). Sub-Saharan Africa’s harsh conditions, with frequent drought and nutrient-poor soil (Sanchez 2002; Tully *et al*. 2015; Connolly-Boutin & Smit 2016), may act as such a strong stressor. In African deserts, deterministic bacterial cooccurrence was uniquely observed, while stochastic processes partially contributed to community assembly in other deserts (Caruso *et al*. 2011). It is possible that environmental selection plays a more prominent role in sub-Saharan Africa than in other regions. Examining microbial community assembly processes in the context of environmental conditions and microbial traits is crucial for assessing the impacts of land-use changes on soil health and devising sustainable management strategies (Allison & Martiny 2008; Dubey *et al*. 2019).

This study compared soil microbial communities across multiple sites in Kenya and Malawi, each with a paired natural ecosystem and farmland. We investigated how microbial taxonomic and functional heterogeneity is influenced by spatial structure, farming activity, and environmental factors, and we clarify how these determinants vary across scales. We hypothesized that farming activities drive the spatial homogenization of microbial functional composition more broadly than that of taxonomic composition. This study contributes to a deeper understanding of how land-use changes affect the taxonomic and functional diversity of soil microbiomes, from within-site (∼200 m) to across-site (∼1500 km) scales.

## Mateirals and Methods

### Site description

In November 2019, soil samples were collected from five sites in Kenya and Malawi, designated as Sites D to H. Each of these sites had adjacent natural and farmed lands (Figure 1). These sites experience average annual rainfall of 1135 mm and 952 mm in Kenya and Malawi, respectively (Data Africa; https://DataAfrica.io, accessed on January 30, 2025). According to local administrators, the farmlands were under cultivation for over 15 years and primarily grew maize, beans, tomatoes, cabbages, and several horticultural crops. Vegetation in the natural lands was generally classified as shrubland (e.g., Acacia bushes) in Kenya and Miombo woodlands in Malawi. Sampling site locations were recorded with a GPS logger (eTrex 20; GARMIN Inc., USA). Based on previous studies and spatial analysis using the Soil Atlas of Africa in QGIS (3.36.3), Sites D and E were classified as Humic Nitisols, Site F as Ferrochromic Luvisols, and Sites G and H as Chromic Luvisols or Haplic Lixisols (Kibunja *et al*. 2010; Dewitte *et al*. 2013; Phiri & Njira 2023). The distance between the two land-use types (natural and farm) was approximately 100 m for each site. For each sampling, three 1 m × 1 m plots, approximately 20 m apart from each other, were set (Table S2). Within each plot, three soil cores (100 cm^3^ each) were sampled, resulting in a total of nine replicates for each treatment. The samples were shipped from Kenya and Malawi to Japan, which took approximately one month. The subsequent experiments were conducted immediately upon their arrival.

**Figure 1.**
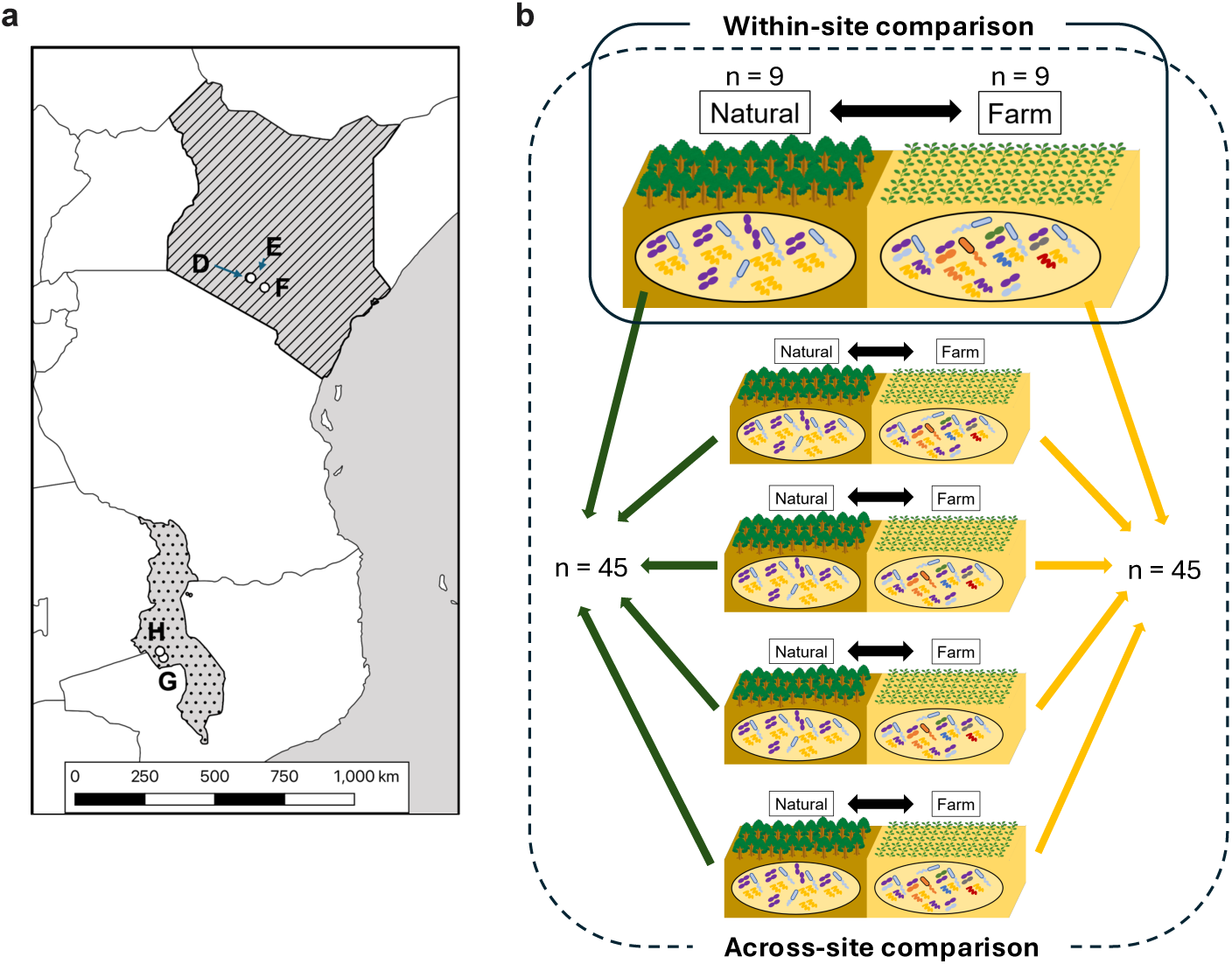
Location of sampling sites and concept of the analysis. (**a**) The sampling locations, Sites D–F in Kenya and Sites G and H in Malawi, are annotated in the map. (**b**) All sites had pairs of neighboring natural (unmanaged) lands and farmlands. Nine soil samples were collected at each land use at one site, resulting in 45 samples for each land use across five sites.

### Measurements of soil physical and chemical properties

Soil pH was determined by mixing 6 g of soil with 30 mL of deionized water, shaking for one hour, and measuring the pH of the supernatant using a pH sensor (AS800; AS ONE Co., Japan). Total C and nitrogen (N) contents were measured using dried soils with an organic elemental analyzer (2400 Series II CHNS/O Elemental Analysis; PerkinElmer Co., USA). Measurements of soil moisture and texture were conducted by researchers at the University of Nairobi and Lilongwe University of Agriculture and Natural Resources. Soil moisture was determined by weighing fresh and dry soils after oven-drying at 105^◦^C for over 24 h. Soil texture was assessed by the separation of sand (> 0.05 mm) with wet sieving of the soil mixture, followed by sampling a suspension with a pipette to remove clay (< 0.002 mm) from the mixture, leaving silt (0.05–0.002 mm). The soil textures are presented in Table S3.

### DNA extraction and quantitative PCR

DNA was extracted from 0.5 g of each soil sample with a soil DNA extraction kit (NucleoSpin Soil; MACHEREY-NAGEL GmbH & Co. KG, Germany). Extraction enhancer SX was used for all samples. Extracted DNA concentrations were determined using a Fluorometer (Quantus, Promega, Madison, WI, USA). Buffer SL1 was initially applied, but samples yielding DNA concentrations below 100 ng/mL were re-extracted using buffer SL2, which provided sufficient DNA for subsequent analyses. Details of samples treated with SL1 or SL2 are summarized in Table S4. We verified that the choice of buffer (SL1 or SL2) did not affect the microbial community structures (Figure S7). Extracted DNA was stored at −30^◦^C until further analysis. The abundances of prokaryotes and fungi were quantified by quantitative PCR using the CFX Connected Real-Time PCR detection system (Bio-Rad, Hercules, CA, USA). The reaction was performed in a 20 *µ*L with 10.4 *µ*L of KAPA SYBR FAST qPCR Master Mix (2X) ROX Low (Kapa Biosystems, Wilmington, MA, USA), 7 *µ*L of nuclease-free water, 0.8 *µ*L of each primer (10 *µ*M), and 1 *µ*L of the 1, 10, 50, 100, or 500-fold diluted DNA sample (Table S4). The prokaryotic 16S ribosomal RNA (16S rRNA) and fungal internal transcribed spacer (ITS) genes were detected using universal primer sets (16S rRNA, 515f: 5^′^-GTGCCAGCMGCCGCGGTAA-3^′^, 806r: 5^′^-GGACTACHVGGGTWTCTAAT-3^′^; ITS, ITS1: 5^′^-TCCGTAGGTGAACCTGCGG-3^′^, ITS2: 5^′^-GCTGCGTTCTTCATCGATGC-3^′^).

The qPCR conditions for 16S rRNA were as follows: initial denaturation at 95^◦^C for 3 min, followed by 35 cycles of 95^◦^C for 30 s, 58^◦^C for 1 min, and 72^◦^C for 1 min. The qPCR conditions for ITS were as follows: initial denaturation at 95^◦^C for 30 s, followed by 35 cycles of 95^◦^C for 30 s, 55^◦^C for 1 min, and 72^◦^C for 1 min. The 16S rRNA and ITS qPCR amplifications were followed by a melting curve analysis. The DNA quantity in the standard 16S rRNA or ITS amplicons from the microbial community standard (ZymoBIOMICS Microbial Community DNA Standard, Zymo Research, Irvine, CA, USA), was determined using Quantus (Promega). The qPCR efficiency was 110%–112% and 76%–80% for 16S rRNA and ITS, respectively.

### DNA amplicon sequencing

The 16S rRNA and ITS genes were amplified by PCR from the extracted DNA and used for further sequencing. The primer sets targeted the same regions as the qPCR described above. For 16S rRNA, the reaction was performed in 20 *µ*L with 10.4 *µ*L of Taq polymerase (AmpliTaq Gold 360 DNA Master Mix, Thermo Fisher Scientific), 8.2 *µ*L of nuclease-free water, 0.4 *µ*L of each primer (10 *µ*M), and 1 *µ*L of diluted DNA samples (Table S4). The PCR conditions for 16S rRNA were as follows: initial denaturation at 95^◦^C for 10 min, followed by 30 cycles of 95^◦^C for 30 s, 57^◦^C for 1 min, and 72^◦^C for 1 min. For ITS, the reaction was performed in 20 *µ*L with 10.4 *µ*L of the AmpliTaq Gold 360 DNA Master Mix (Thermo Fisher Scientific), 8.02 *µ*L of nuclease-free water, 0.4 *µ*L of each primer (10 *µ*M), 0.18 *µ*L of bovine serum albumin, and 1 *µ*L of diluted DNA samples (Table S4). The PCR conditions for ITS were as follows: initial denaturation at 95^◦^C for 10 min, followed by 30 cycles of 95^◦^C for 30 s, 53^◦^C for 15 s, and 72^◦^C for 30 s. After the PCR reactions, the samples were purified using a DNA purification kit (AMPure XP Kit; Beckman Coulter, Inc., USA) to remove primers and enzymes. The amplified DNA was labeled with Ion Xpress barcode adapters (Thermo Fisher Scientific) by a PCR of five cycles in the same conditions as described above (except for the number of cycles). After purification with the AMPure, the molarity of the products was quantified using the Bioanalyzer High Sensitivity DNA Kit (Agilent Technologies, USA). The samples were then mixed and diluted to 50 pM and loaded into an Ion 318 Chip, using the Ion Chef Instruments (Thermo Fisher Scientific) with the Ion PGM™Hi-Q™View Chef Kit. The Ion PGM™Hi-Q™View Sequencing Kit and the Ion PGM™Sequencer (Thermo Fisher Scientific) were used for DNA sequencing.

### Analysis of sequence data and annotation of predicted functions and traits

The sequence data was obtained from the Ion Torrent system and analyzed using the dada2 package on R (4.3.3) for adapter trimming, denoising, chimera removal, and determination of amplicon sequencing variant (ASV) (Callahan *et al*. 2016). For the adapter trimming, the sequences were truncated by detecting the 16S rRNA and ITS primers using the “cutadapt” command (Martin 2011). Taxonomic information of the obtained ASVs was annotated using the assignTaxonomy function in R, referring to databases SILVA (release 132) for 16S rRNA and UNITE (8.3) for ITS (Quast *et al*. 2013; Nilsson *et al*. 2019). The mitochondria and chloroplasts comprised in SILVA were removed from the prokaryotic ASV table, as described previously (Tatsumi *et al*. 2023b). We normalized ASV counts across samples using rarefaction based on the minimum read number, equalizing the prokaryotic and fungal reads in each sample to 10,193 and 7,564, respectively. In total, 9,425 and 9,074 unique ASVs were found across samples for 16S rRNA and ITS, respectively. The Shannon diversity index for each sample was calculated using the diversity function the vegan package in R. A PICRUSt2 analysis was performed on the prokaryotic taxa to predict prokaryotic functional genes (Douglas *et al*. 2020). From all predicted functions, C cycle-related and N cycle-related functions were selected based on KEGG Orthology (Kanehisa *et al*. 2016), referring to previous studies (Kaiser *et al*. 2016; Alfreider *et al*. 2018; Zhang *et al*. 2020a) (Table S5). The prokaryotic C cycleand N cycle-related functions were categorized into process categories: “Autotroph,” “Cellulolytic,” “Chitinolytic,” “Lignolytic,” “Xylanolytic,” “Methanotroph,” “Methanogen,” “N-fixing,” “Nitrifying,” and “Denitrifying.” Fungal taxa were searched against the FungalTraits database to assign fungal traits (Põlme *et al*. 2020). The “primary_lifestyles” in FungalTraits was used for the following analyses.

### Microbial community dissimilarity analysis

The dissimilarities in the prokaryotic and fungal community structures were analyzed through non-metric multidimensional scaling using the metaMDS function from the vegan package (Dixon 2003) in R. Permutational multivariate analysis of variance (PERMANOVA) was employed to assess the differences in the community structures between the land-use types and among sites. This analysis considered the factors “land use” (Natural or Farm) and “site” (Sites D, E, F, G, or H) using the adonis2 function from the vegan package and the “bray” method with 100,000 permutations. The linear fit between the microbial communities and the prokaryotic functions, fungal lifestyles, and environmental factors was tested by 100,000 permutations using the envfit function in the vegan package. For the calculation, the C and N cycle-related functions as described earlier were used for prokaryotic functions, while “Pathotroph,” “Saprotroph,” and “Ectomycorrhizal” fungal lifestyles were selected based on the primary lifestyle classification in FungalTraits. The ANCOMBC test was conducted using the ancombc function in R to examine differences in the relative abundances of microbes at the class level between land uses (Lin & Peddada 2020).

Microbial community heterogeneity was assessed by calculating dispersion (i.e., distances to group centroids) within each treatment (i.e., Site D–Natural, Site D–Farm, Site E–Natural, . . ., Site H–Farm) and for each land-use type across sites (i.e., Natural or Farm across all sites).

The calculations were performed using the “bray” method with the vegdist and betadisper functions in R. Additionally, the heterogeneity of all prokaryotic functions, C and N cycle-related functions, fungal primary lifestyles, and environmental factors were calculated using the same method.

Autocorrelations in taxonomic and functional compositions were quantified using Mantel’s test with the mantel.correlog function in R. Distance classes were defined as within-site (0– 200 m), across sites within the country (200–30,000 m and 30,000–70,000 m), and across sites between countries (i.e., between Kenya and Malawi; 70,000–1,444,000 m and 1,444,000– 1,500,000 m).

The contributions of each fungal ASV to the land-use group heterogeneity were quantified at three spatial scales: “within site,” “across sites within the country,” and “across sites between the countries,” using similarity percentage analysis (Clarke 1993). First, the contributions of each ASV to Bray–Curtis dissimilarity was calculated in all sample pairs on a geographic scale with the simper function of the vegan package in R. The average contribution within each scale was then computed. ASVs with an average contribution greater than 0 were grouped by their primary lifestyles. Differences in the average contributions of each lifestyle between land uses were tested using a permutation test (1,000 iterations), shuffling samples within each site while not preserving land-use labels (Natural or Farm).

### Community assembly processes and variation partitioning

Ecological processes of community assembly were estimated using the *β*-nearest taxon index (*β*NTI) and Raup–Crick based on Bray–Curtis dissimilarity (RC_bray_) following previous studies (Stegen *et al*. 2012; Peng *et al*. 2024). We quantified *β*NTI for all sample pairs for prokaryotes and fungi using ASV-level composition and phylogenetic trees. Phylogenetic trees were constructed using the Neighbor-Joining method with the phangorn package in R. A |*β*NTI| value > 2 indicates that community assembly primarily driven by selection, whereas |*β*NTI| < 2 suggests that assembly is influenced by drift, dispersal limitation, and homogenizing dispersal. More specifically, *β*NTI < −2 represents significantly less phylogenetic turnover than expected under null models (homogeneous selection), while *β*NTI > 2 indicates significantly greater phylogenetic turnover than expected (variable selection). For pairs where |*β*NTI| < 2, RC_bray_ was used to further differentiate assembly processes: homogenizing dispersal (|*β*NTI| < 2 and RC_bray_ < −0.95), dispersal limitation (|*β*NTI| < 2 and RC_bray_ > 0.95), and undominated processes (|*β*NTI| < 2 and |RC_bray_| < 0.95), reflecting weak selection, weak dispersal, diversification, or drift.

Variation partitioning analysis was conducted using the varpart function in R to assess the effects of spatial distance, environmental factors, their overlaps, and unexplained variations on taxonomic and functional compositions. Moran’s Eigenvector maps (MEMs) were used to model spatial variation (Dray *et al*. 2006). Candidate spatial weight matrices were generated with Gabriel and Relative Neighborhood Graphs using binary and linear weighting schemes. The optimal weighting procedure was selected based on its ability to capture spatial patterns through 99 permutations.

### Statistics

A two-way analysis of variance (ANOVA) was performed to assess the variations in soil C and N contents, pH, water content, 16S rRNA and ITS copy numbers, and Shannon diversity indices of prokaryotic and fungal communities, considering the land use (Natural or Farm) and site (Sites D, E, F, G, or H) factors. Normality and homogeneity of variances were evaluated using the Shapiro–Wilk test in R. If p-values were below 0.05, data were square-root transformed before performing the two-way ANOVA. If a significant interaction between land use and site was detected in the two-way ANOVA, pairwise comparisons within each site were conducted using the emmeans function in R. Distance to group centroids for within-site heterogeneity was tested using a two-way ANOVA following the same procedure, while across-site heterogeneity was assessed using a t-test. All correlations in this study were tested by Pearson’s method using the cor.test function in R. Factors driving microbial homogenization were identified through a multiple regression analysis, employing soil physicochemical properties (farming, moisture, total C, total N, and pH) using the lm and tidy functions in R.

## Results

### Soil characteristics

Effects of land use and site on soil properties are presented in Table 1. The soil C content was 29.7%, 28.2%, 32.0%, and 31.3% lower in farmlands than in counterpart natural lands at Sites D, F, G, and H, respectively. The total N content was 31.3% lower in farmland than in natural land at Site D. The C/N ratio was 14.4% lower in farmlands than in natural lands when averaged across sites (p < 0.05). The soil pH was significantly lower in farmlands than in natural lands at all sites except Site E. The prokaryotic gene (16S rRNA) abundance was 8.8% lower in farmlands than in natural lands when averaged across sites (p < 0.001). The fungal gene (ITS) abundance was 13.9%, 19.9%, and 9.7% lower in farmed soil than in natural soil at Sites D, G, and H, respectively. The effect of land use on the Shannon diversity index of prokaryotes and fungi varied among sites (Table 1).

**Table 1.** Soil biogeochemical properties.

| Characteristic | Site D |  | Site E |  | Site F |  | Site G |  | Site H |  | Two-way ANOVA p-values |  |  |
| --- | --- | --- | --- | --- | --- | --- | --- | --- | --- | --- | --- | --- | --- |
|  | Natural<sup>1</sup> | Farm<sup>1</sup> | Natural<sup>1</sup> | Farm<sup>1</sup> | Natural<sup>1</sup> | Farm<sup>1</sup> | Natural<sup>1</sup> | Farm<sup>1</sup> | Natural<sup>1</sup> | Farm<sup>1</sup> | Site | Landuse | Site × Landuse |
| Moisture (%) | **45.4 ± 2.7** | **35.4 ± 1.1** | 33.8 ± 2.3 | 31.7 ± 1.8 | 19.0 ± 4.5 | 18.5 ± 2.0 | **4.9 ± 0.9** | **3.7 ± 1.1** | **4.6 ± 1.8** | **2.6 ± 0.6** | <0.001 | <0.001 | 0.002 |
| Total C (%) | **3.1 ± 0.7** | **2.2 ± 0.3** | 2.0 ± 0.3 | 2.2 ± 0.2 | **1.3 ± 0.6** | **1.0 ± 0.5** | **1.3 ± 0.2** | **0.9 ± 0.1** | **2.8 ± 0.5** | **1.9 ± 0.3** | <0.001 | <0.001 | 0.004 |
| Total N (%) | **0.4 ± 0.1** | **0.3 ± 0.1** | 0.3 ± 0.1 | 0.4 ± 0.1 | 0.3 ± 0.1 | 0.3 ± 0.1 | 0.2 ± 0.1 | 0.2 ± 0.1 | 0.3 ± 0.1 | 0.2 ± 0.0 | <0.001 | 0.3 | 0.030 |
| C/N ratio | **9.1 ± 2.3** | **9.5 ± 3.2** | **7.4 ± 3.0** | **6.3 ± 1.3** | **4.1 ± 1.7** | **3.2 ± 2.3** | **10.4 ± 4.8** | **7.6 ± 7.6** | **10.7 ± 2.0** | **9.0 ± 2.4** | <0.001 | 0.040 | 0.5 |
| pH (H<sub>2</sub>O) | **6.6 ± 0.1** | **6.1 ± 0.1** | 6.6 ± 0.2 | 6.7 ± 0.3 | **6.7 ± 0.1** | **5.4 ± 0.4** | **7.2 ± 0.2** | **5.8 ± 0.1** | **6.2 ± 0.1** | **5.6 ± 0.1** | <0.001 | <0.001 | <0.001 |
| 16S rRNA (log copies g<sup>-1</sup>soil) | **8.3 ± 0.6** | **7.4 ± 1.1** | **7.6 ± 1.1** | **7.4 ± 1.0** | **8.2 ± 0.4** | **7.5 ± 0.5** | **8.3 ± 0.2** | **7.1 ± 0.6** | **8.1 ± 0.4** | **7.5 ± 0.5** | 0.5 | <0.001 | 0.2 |
| ITS (log copies g<sup>-1</sup>soil) | **7.5 ± 0.6** | **6.4 ± 1.0** | 6.6 ± 1.1 | 6.5 ± 1.0 | 7.4 ± 0.2 | 6.9 ± 0.5 | **7.8 ± 0.3** | **6.2 ± 0.6** | **7.5 ± 0.3** | **6.7 ± 0.5** | 0.13 | <0.001 | 0.038 |
| 16S rRNA (Shannon Diversity) | **7.2 ± 0.3** | **7.6 ± 0.1** | 7.5 ± 0.2 | 7.4 ± 0.3 | **7.4 ± 0.4** | **8.4 ± 0.6** | **8.4 ± 0.1** | **8.0 ± 0.4** | 8.4 ± 0.3 | 8.3 ± 0.2 | <0.001 | 0.023 | <0.001 |
| ITS (Shannon Diversity) | **4.6 ± 0.9** | **5.6 ± 0.8** | 6.0 ± 0.3 | 6.0 ± 0.2 | 6.0 ± 0.7 | 5.5 ± 0.3 | 5.5 ± 0.4 | 5.1 ± 0.5 | **5.3 ± 0.9** | **6.0 ± 0.3** | <0.001 | 0.2 | <0.001 |

<sup>1</sup>Mean ± SD in each location (n=9). Mean values significantly different between land uses, as determined by two-way ANOVA or posthoc test, are shown in bold.

### Microbial community structures, functional characteristics, and their taxonomic heterogeneity

In PERMANOVA analysis, there was an interaction between land use and site for both prokaryotic and fungal communities, but clear separations by land use were observed (Figure 2a, b). As for relative abundances at the class level between land uses, the prokaryotes Ktedonobacteria, Bacilli, Chthonomonadetes, JG30-KF-CM66, Armatimonadia, Acidobacteriia, and OLB14 and the fungi Sordariomycetes, Rhizophlyctidomycetes, GS13, and Dothideomycetes exhibited significantly larger abundances in farmlands when averaged across sites (Figure S1a, b). For the prokaryotic communities, various C and N cycle-related functions were positively correlated with communities in Sites F, G, and H, with stronger correlations observed in natural lands (Figure 2a). The methanotroph and N-fixing functions tended to be more abundant in farmland communities, particularly at Site F (Figure 2a). The nitrifying function was predominantly observed in communities in Sites D and E, to which the total N content was also positively correlated (Figure 2a). For the fungal communities, saprotrophs were more abundant in the natural land communities, while pathotrophs were prominent in the farmland communities (Figure 2b). The heterogeneity of prokaryotic communities within each site, assessed by distances to centroids, was significantly smaller in farmed soils than in natural soils (Figure 2c). An interaction between site and land use in the heterogeneity of fungal communities was observed, and the heterogeneity was significantly smaller in farmland at Site G (Figure 2d). At the across-site scale, no significant difference in prokaryotic heterogeneity between land uses was observed, whereas fungal heterogeneity was significantly smaller in farmlands (Figure 2e, f).

**Figure 2.**
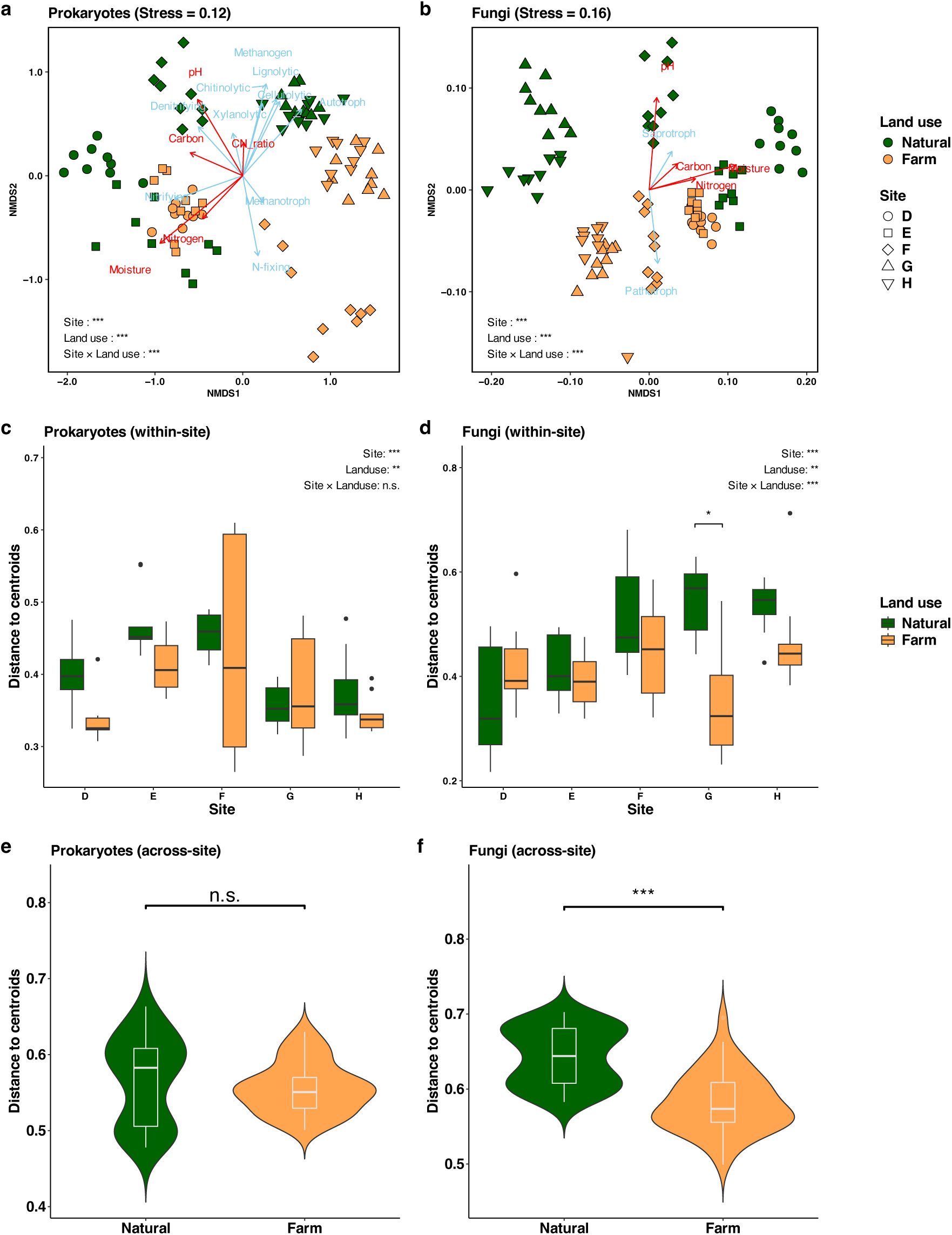
Dissimilarity of soil microbial communities in natural lands and farmlands. NMDS for (**a**) prokaryotes and (**b**) fungi are shown. Prokaryotic or fungal functions and environmental variables that significantly correlated with the communities are illustrated with arrows, colored by red and skyblue, respectively. Distance to centroids of the communities in each land use in (**c, d**) within-site and (**e, f**) across-site scales are plotted, for prokaryotes (**c, e**) and fungi (**d, f**). The p-values in the PERMANOVA or two-way ANOVA on sites, land uses and their interaction, and those in the t-test on land uses are indicated with “*”, “**”, or “***”, representing p < 0.05, p < 0.01, or p < 0.001, respectively. If a significant interaction was found in the two-way ANOVA, pairwise comparisons of estimated marginal means were conducted to assess whether there were significant differences in land use within each site. Significant differences in land use within sites were indicated by a single asterisk “*”.

### Relationships between taxonomic homogenization and functional homogenization

At the within-site scale, positive correlations were observed between the heterogeneities of prokaryotic taxa and their functional composition at all sites (Figure 3a). Similarly, fungal taxonomic heterogeneity was positively correlated with the heterogeneity of their lifestyles at all sites except Site G (Figure 3b). At the across-site scale, no significant correlation was observed between the heterogeneities of prokaryotic taxa and their functions, whereas fungal taxonomic heterogeneity showed a positive correlation with lifestyle heterogeneity (Figure 3c, d).

**Figure 3.**
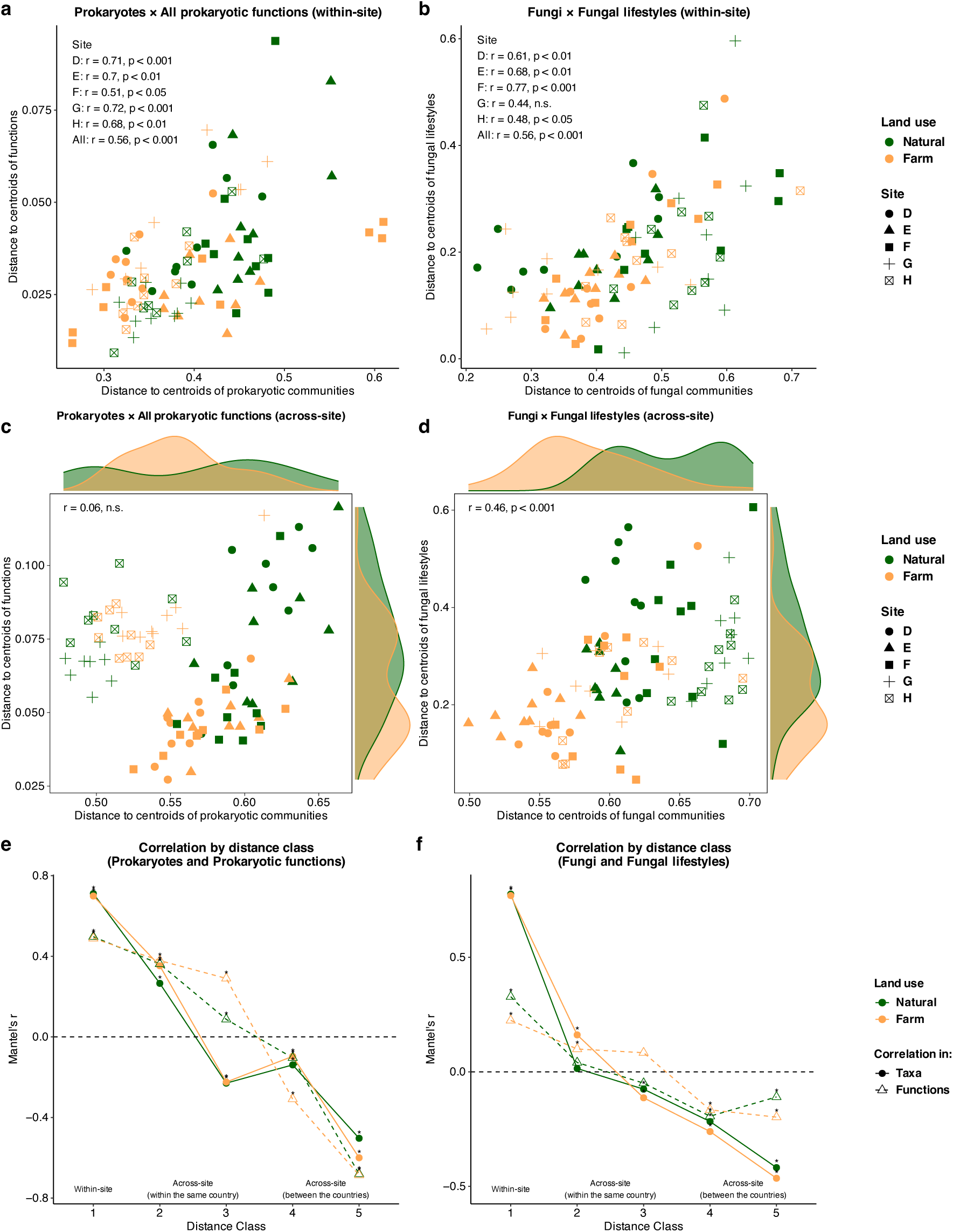
Relationships between heterogeneities of microbial communities and their functions. Correlations between distance to centroids of the microbial communities and that of their functions in each land use within site for (**a**) prokaryotes and (**b**) fungi, and across sites for (**c**) prokaryotes and (**d**) fungi are shown. The correlation coefficients and p-values in the Pearson’s correlation tests are indicated. Mantel’s correlograms of microbial community compositions and functional compositions by distance class are plotted for (**e**) prokaryotes and prokaryotic functions and (**f**) fungi and fungal lifestyles. The significant autocorrelations (corrected-p < 0.05) in each distance class tested by permutation is indicated with *.

The Mantel correlograms revealed that the autocorrelations of prokaryotic and fungal taxa at Distance class 2 (i.e., 200–30,000 m) were higher in farmlands than in natural lands, indicating microbial taxonomic homogenization among farmlands at this spatial scale (Figure 3e, f). Notably, functional autocorrelations retained higher in farmlands compared to natural lands even at Distance class 3 (i.e., 30,000–70,000 m), where taxonomic autocorrelations were no longer observed (Figure 3e, f).

### Community assembly processes and partitioning of taxonomic and functional variations by spatial and environmental factors

Across all spatial scales and land uses, the community assembly process of prokaryotes was predominantly governed by homogeneous selection (Figure 4a). In contrast, the fungal community assembly process varied depending on the scale: homogenizing dispersal was observed primarily at the within-site scale, while dispersal limitation was more prominent at the across-site scales (Figure 4a). Furthermore, dispersal limitation was relatively more dominant in farmland fungal communities compared to natural land fungal communities at the within-site scale, whereas the opposite trend was observed at the across-site scales (Figure 4a). Additionally, across all scales, the contribution of the undominated processes was consistently lower in farmland fungal communities compared to natural land fungal communities (Figure 4a).

**Figure 4.**
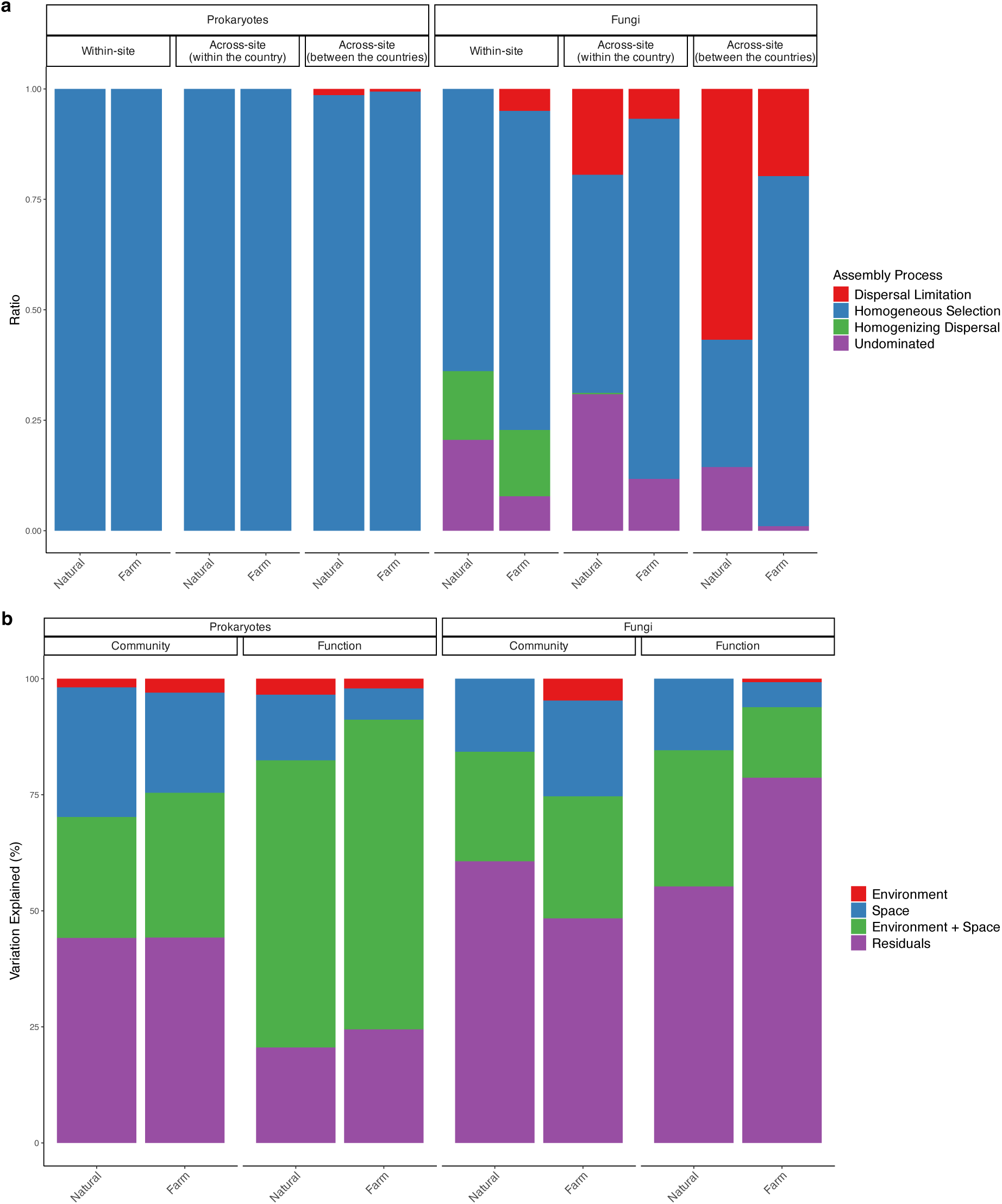
Community assembly process and drivers of taxonomic and functional variation. (**a**) The community assembly process estimated by *β*NTI and RC_bray_ was grouped by the within-site, across-site (within the country), and across-site (between the countries) scales. (**b**) The variations in taxonomic and functional compositions that were explained by environmental factors, space, their overlap, and residuals were shown.

The variation in both prokaryotic and fungal taxa was more strongly explained by “Environment” and “Environment + Space” in farmlands than in natural lands (Figure 4b). Meanwhile, the variation in fungal taxa explained by “Residuals” was relatively lower in farmlands than in natural lands (Figure 4b). For functional compositions, the variations in both prokaryotic and fungal functions in farmlands were less explained by “Space” and had higher “Residuals” compared to functional variation in natural lands.

### Environmental factors related to homogenization of microbial communities

At the within-site scale, prokaryotic community heterogeneity was negatively affected by total C but positively influenced by total N and pH deviation (Figure 5a). In contrast, fungal community heterogeneity was negatively affected by soil moisture, while pH and pH deviation had positive effects (Figure 5b). At the across-site scale, soil moisture had opposing effects on prokaryotic and fungal heterogeneity: positive for prokaryotes and negative for fungi (Figure 5c, d). Additionally, farming activity negatively impacted the heterogeneity of both prokaryotic and fungal communities at the across-site scale (Figure 5c, d).

**Figure 5.**
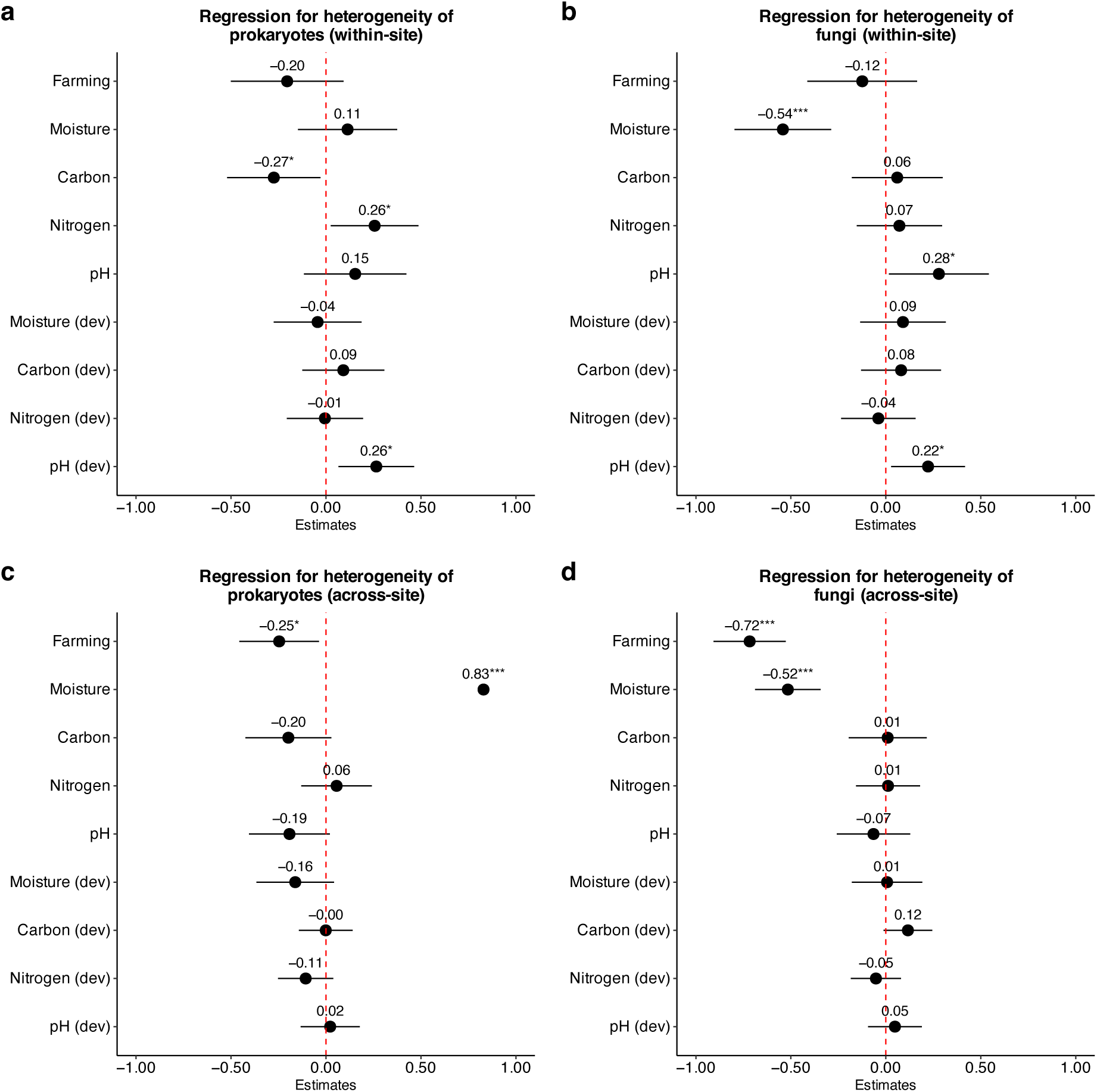
Contributions of farming activity and environmental variables to the heterogeneity of microbial community. The 95% confidence intervals of the regression coefficients, calculated in the multiple regression from farming activity, environmental variables, and absolute deviations of environmental variables to the distance to centroids of (**a, c**) prokaryotes and (**b, d**) fungi in the (**a, b**) within-site and (**c, d**) across-site scales, are shown. The values of coefficients and significance as indicated with “*”, “**”, or “***”, representing p < 0.05, p < 0.01, or p < 0.001, respectively, are noted above each point.

### Relationships between the heterogeneity of fungal communities and fungal lifestyles

The heterogeneity of the fungal community was negatively correlated with the relative abundance of pathotrophs at three sites at the within-site scale (Figure 6a). This trend was even more pronounced at the across-site scale (Figure 6b). Fungal lifestyles contributing to heterogeneity within the farmland group, compared to the natural land group, were mostly plant pathogens (the majority of pathotrophs; Figure S5a) and soil saprotrophs (Figure 6c). In contrast, the contributions of animal parasites and soil saprotrophs were significantly lower in the farmland group at the across-site scale (Figure 6c). The greater contribution of plant pathogens to heterogeneity within the farmland group was consistent across all spatial scales (Figure S4).

**Figure 6.**
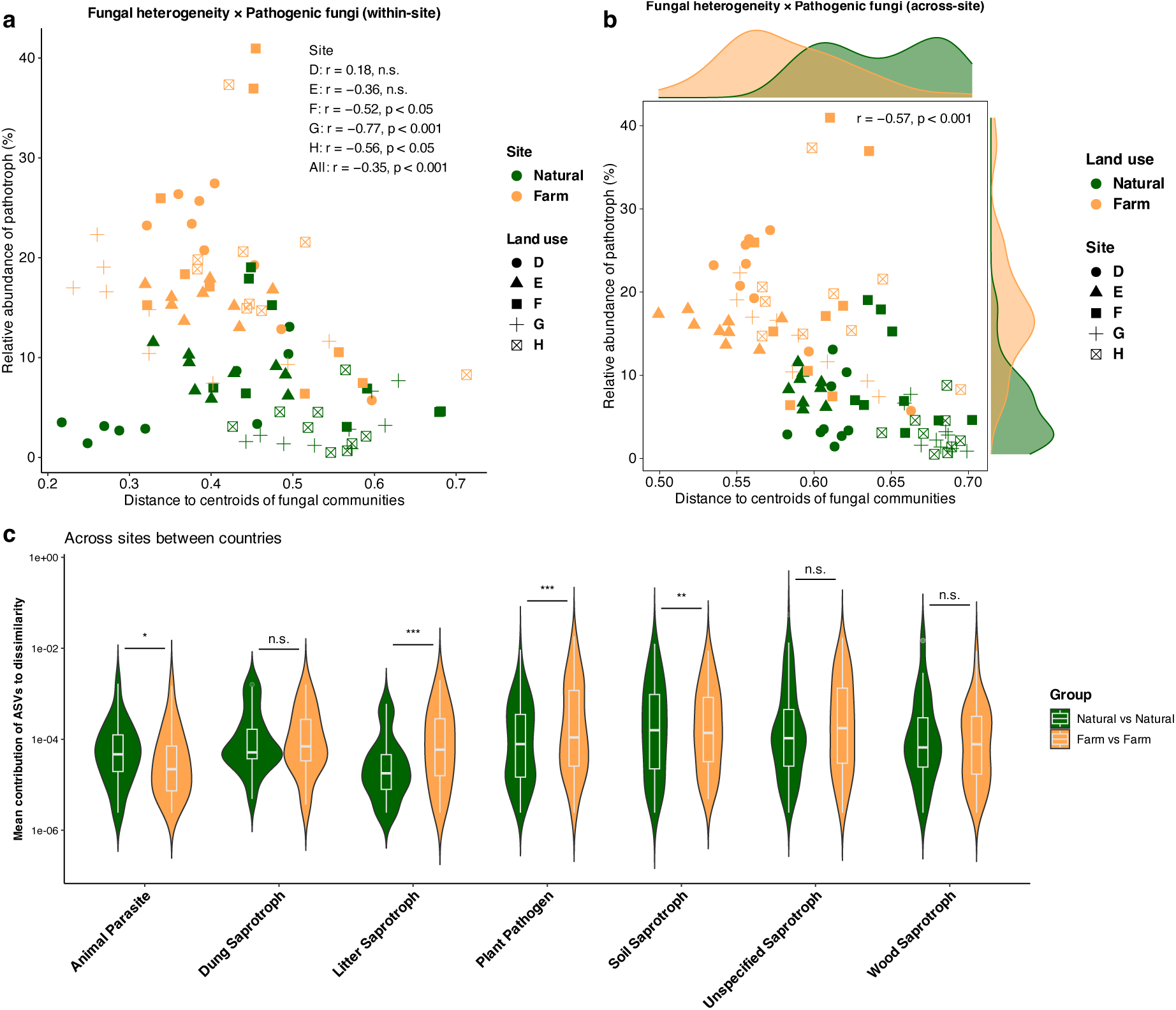
Relation between the heterogeneity of fungal communities and abundance of pathogenic fungi. Correlations between distance to centroids of the fungal communities and relative abundance of pathogenic fungi are plotted for the (**a**) within-site and (**b**) across-site scales. The correlation coefficients and p-values in the Pearson’s correlation tests are indicated. (**c**) The contributions of each fungal ASV to the Bray–Curtis dissimilarity among samples within each land use across site between countries were averaged and grouped by fungal lifestyle. Asterisks (“*”, “**”, or “***”) and n.s. indicate the significance levels of adjusted p-values: < 0.05, p < 0.01, p < 0.001, or no significant difference respectively, tested by 1000 permutations to assess differences in the mean values between natural lands and farmlands.

## Discussion

### Broader homogenization of functions than taxa in soil microbial communities

Our study revealed the relationship between farming-induced taxonomic and functional homogenizations. The positive correlations between taxonomic heterogeneity and functional heterogeneity at the within-site scale, except for one site for fungi, suggest that farming-induced local taxonomic homogenization could also homogenize the functional composition (Figures 2c, d and 3a, b). This observation aligns with a comparable previous study that showed a positive correlation between taxonomic and functional dissimilarity of soil bacteria in the Tibetan Plateau (Liang *et al*. 2023). Contrastingly, at the across-site scale, we observed no clear correlation between the heterogeneity of prokaryotic taxa and their functions, whereas the heterogeneity of fungal taxa was positively correlated with that of their functions (Figure 3c, d). This result may be due to the heterogeneity of prokaryotic taxa varying across sites, particularly between the two countries, whereas the heterogeneity of prokaryotic functions varied with land use (Figures 3c and S2c). Furthermore, even at the scale where prokaryotic taxa lost positive autocorrelation in both land uses (i.e., Distance class 3), their functions retained higher positive autocorrelation in farmlands compared to natural lands (Figure 3e). A similar trend was observed in farmland fungal functions, although the positive autocorrelation was not statistically significant (p = 0.12; Figure 3f). These results suggest that microbial functional compositions among separate farmlands are homogenized even at scales where their taxonomic compositions are dissimilar. Even among different taxa, microbes can perform the same ecosystem functions (Louca *et al*. 2018). Previous studies have revealed that, under identical conditions, bacterial and fungal communities from different sources exhibited functional similarity rather than taxonomic compositional similarity (Langenheder *et al*. 2005, 2006; Knight *et al*. 2024). These findings indicate that the experiment conditions act as selective pressures on microbial functions. Similarly, farming activities homogenize landscapes across different sites (Ribeiro *et al*. 2016), requiring soil microbes to express similar functions. Therefore, it is reasonable to consider that farming activities in our study exerted consistent selecting pressures, driving the homogenization of prokaryotic and fungal functionality over taxonomy. Like other organisms, microbes face barriers to the widespread distribution of certain taxa due to environmental variations, limitations in dispersal ability, and historical factors (Peay *et al*. 2010; Hanson *et al*. 2012; Li *et al*. 2020). However, their functions can surpass these barriers.

### Scale di**ff**erences between prokaryotes and fungi in community homogenization

The homogenization of soil microbial communities due to farming practices occurred at different scales for the major microbial groups; prokaryotic communities were homogenized within each farmland, while fungal communities were homogenized across multiple farmlands (Figure 2c–f). Nearly all prokaryotic clades, at sub-species resolutions (e.g., ≤ 1% average nucleotide difference), are estimated to be globally distributed (Louca 2022). Our results support this estimation, revealing a higher autocorrelation in prokaryotic communities compared to fungal communities (i.e., ∼Distance class 2) and a small degree of dispersal limitation even at broad spatial scales, regardless of land use (Figures 3e, f and 4a). In other words, prokaryotic communities inherently retain a high level of community similarity among different sites. However, prokaryotic community structures are also sensitive to microscale environmental factors, including pH, nutrient availability, and other localized conditions (Lauber *et al*. 2008; Kuramae *et al*. 2012; Delgado-Baquerizo *et al*.2017). Thus, site-specific environmental changes due to farming may locally shape prokaryotic community structures, leading to the within-site homogenization observed in this study (Figure 2c).

In contrast, fungal communities exhibited lower autocorrelation and were more influenced by dispersal limitation than prokaryotes, indicating that they maintain relatively greater variation among sites (Figures 3e, f and 4a). The steeper distance-decay of community similarity or greater dispersal limitation in the fungal community, compared to prokaryotes, agrees with previous studies (Luan *et al*. 2020; Zhang *et al*. 2020b; Zhang *et al*. 2021). Our study, however, also demonstrated that farming activity reduced dispersal limitation in fungal communities across different farmlands, leading to across-site homogenization (Figures 2f and 4a). This effect may be attributed to the spore-related dispersal abilities of certain fungi, which enable relatively efficient long-distance spread (Tedersoo *et al*. 2014). Indeed, in our study, the fungal genera that displayed greater relative abundance in farmlands included *Fusarium*, *Aspergillus*, and *Alternaria* (Figure S1d), which are known for their long-distance dispersal potential (Golan & Pringle 2017). Hence, while fungal communities originally varied among sites, farming may create specific niches for some opportunistic species simultaneously across different sites, driving community homogenization at broad spatial scales.

### Factors driving microbial homogenizations

The heterogeneity of composite environmental factors showed no correlation with the heterogeneity of microbial communities at any scale (Figure S3). Hence, in sub-Saharan Africa, the environmental heterogeneity that arose from various factors was secondary; rather, each environmental factor or its variation may solely influence the heterogeneity of microbial communities (Figure 5). By contrast, as demonstrated in several studies, the heterogeneous composition of environmental factors seems to enhance the heterogeneity of soil microbial communities in other regions, including North America and China (Ramette & Tiedje 2007; Liu *et al*. 2023; Tatsumi *et al*. 2023a). The strong association between individual environmental factors and microbial heterogeneity in our study may be due to the relatively stressed conditions in sub-Saharan Africa, such as drought stress (Connolly-Boutin & Smit 2016), which increases the importance of environmental selections in community assembly (Ning *et al*. 2024). The dominance of homogeneous selection in the community assembly process indicates consistent environmental selection throughout this region (Figure 4a).

The individual factors contributing to the heterogeneity of microbial communities varied depending on the scales (Figure 5). At the within-site scale, several environmental factors, rather than farming itself, influenced the microbial heterogeneity (Figure 5a, b). As a common tendency for prokaryotes and fungi, higher soil pH or pH variation induced their heterogeneity (Figure 5a, b). This observation is consistent with many studies describing niche specialization and community changes in soil microbes with pH (Dumbrell *et al*. 2010; Rousk *et al*. 2010; Prosser & Nicol 2012; Tripathi *et al*. 2018; Duan *et al*. 2023). Additionally, as unique effects for prokaryotic heterogeneity on the within-site scale, soil N had a positive influence, whereas soil C had a negative one (Figure 5a). In C-rich environments with limited N and phosphorus amounts, competitive species tend to dominate, leading to lower species *α* diversity (Tardy *et al*. 2015; Delgado-Baquerizo *et al*. 2017; Ohigashi *et al*. 2021). In this study, prokaryotic communities with high C-degrading functions (Cellulolytic, Lignolytic, Chitinolytic, and Xylanolytic) showed strong positive correlations with the C/N ratio (Figure 2a). This finding suggests that C-degrading species may gain a competitive advantage in such C-rich but N-poor environments, possibly resulting in the formation of similar communities. In contrast, moderate N availability can promote prokaryotic community heterogeneity, as observed in a long-term N fertilization experiment (Liu *et al*. 2021). This effect is likely due to a strengthened priority effect with increased nutrient availability, where the order and timing of species’ arrival have a bigger influence on community assembly (Fukami 2015; Liu *et al*. 2021). Although both C and N are essential nutrients for prokaryotic growth, a previous study showed that a low C/N ratio promotes an increase of copiotrophs, organisms with high maximum growth rates, necessitating consideration of the balance of these nutrients in community assembly (Wang *et al*. 2024). Therefore, soil N, but not C, may provide early-arriving copiotrophs with opportunities to persist, contributing to prokaryotic community heterogeneity within a site (Figure 5a).

As for fungal communities, soil moisture negatively affected within-site heterogeneity (Figure 5b). Soil fungi tend to thrive in large soil pores, which fill with water when moisture levels are high, while prokaryotes can inhabit small pores, where they are more protected from such perturbations (Kaisermann *et al*. 2015). This suggests that soil moisture may have acted as a selective pressure for fungal communities, reducing their heterogeneity and contributing to their homogenization (Figure 5b).

On the other hand, at the across-site scale, farming itself was crucial in microbial homogenization (Figure 5c, d). This observation indicates that “farming activity,” which is unexplained by a specific environmental factor but homogenizes landscapes at various locations, indeed plays a key role in homogenizing soil microbial communities across the locations. This result agrees with previous studies documenting farming-induced microbial homogenization in China and South America and global meta-analysis (Rodrigues *et al*. 2013; Peng *et al*. 2024). Conversely, some studies demonstrated that croplands, including non-till farmlands, retained higher heterogeneity of soil bacteria than natural steppe or managed grasslands (Goss-Souza *et al*. 2017; Liu *et al*. 2023). These results may seem contradictory, but they are consistent when considering that the more homogenized landscapes, rather than limiting within “farmlands,” likely lead to microbial homogenization. Goss-Souza et al. (2017) suggested that high dispersal and selection rates were driving bacterial community homogenization in grasslands, a pattern that aligns with the processes observed in farmlands in our study (Figure 4a).

In contrast to the negative effect of farming, soil moisture positively affected the across-site prokaryotic heterogeneity (Figure 5c). This observation may seem to contradict the fact that soil moisture connects soil pores and facilitates prokaryote movement, leading to local homogenization (Carson *et al*.2010; Bickel & Or 2020). However, such movement would not occur across sites in the current study (∼1,500 km). The increased mobility of prokaryotes at the microscale due to soil moisture may facilitate stochastic assembly processes when comparing distant sites (Farjalla *et al*. 2012; Luan *et al*. 2020), potentially contributing to across-site heterogeneity. This explanation is supported by the observation that relatively stochastic prokaryotic assembly, indicated by absolute *β*NTI approaching lower values, occurred at intermediate moisture levels between corresponding soil pair (Figure S6a). In contrast, fungi grow hyphae and do not frequently move through soil pores (Tecon & Or 2017), which may explain why increased soil moisture does not promote a drastic turnover from deterministic to stochastic processes, unlike prokaryotes, even under higher soil moisture conditions (Figure S6b). As a result, the vulnerability to moisture, as described above, may still act as a selective pressure to homogenize fungal communities (Figure 5d).

### Microbial community homogenization and ecosystem functions

Our study revealed that the microbial functional composition was more homogenized in farmlands than in natural lands (Figures 3 and S2). The prokaryotic N-fixing and methanotrophic functions, strongly correlated with farmland communities, may represent the homogenized functions (Figure 2a). These functions provide N to soil and oxidize methane (a greenhouse gas), respectively, and are indicators of ecosystem services (Lew & Glińska-Lewczuk 2018; Schloter *et al*. 2018). Thus, farming-induced functional homogenization may sometimes result in an enhancement of certain “profitable functions.”

However, such imbalanced functionality alters key soil processes and can further drive pathogen emergence (Eisenhauer *et al*. 2013; Trivedi *et al*. 2019). In our study, more abundant pathotrophs were observed in the more homogenized fungal communities (Figure 6a, b), leading us to consider that increased pathotrophs themselves may be a defining feature of the observed homogeneity. However, pathotrophic ASVs did not directly contribute to the fungal community homogenization in farmlands; rather, they contributed to maintaining heterogeneity (Figure 6c). This discrepancy suggests two possible scenarios: (1) fungal community homogenization in farmlands indirectly facilitated the relative increase in pathotrophs, or (2) fungal community homogenization was not associated with the relative increase in pathotrophs in farmlands. Scenario (1) may occur as diverse communities and functional dissimilarity can protect against invader species by efficiently utilizing resources, thereby reducing the resource availability for the invaders (Eisenhauer *et al*. 2013; Bonanomi *et al*. 2014; Mallon *et al*. 2015). The lower dispersal limitation observed in fungal communities in farmlands at the across-site scale may indicate fewer barriers to the widespread distribution of invaders (Figure 4a). However, scenario (2) is still possible because some pathogenic fungi grow in acidic soils with pH < 6.5 (Zhang *et al*. 2022), which are similar to the conditions in the farmlands of our study (Table 1). The negative correlation between soil pH and the relative abundance of several pathogenic fungal genera supports this scenario (Table S1). Additionally, monoculture decreases the competition between fungal species, allowing pathogenic fungi to increase (Wang *et al*. 2023). Since our studied farmlands grew few plant species, including maize, and the farming-induced genera *Fusarium*, *Epicoccum*, and *Curvularia* infect maize (Oldenburg *et al*. 2017; Manzar *et al*. 2021; Xu *et al*. 2022), crop management may also play a role (Figure S1). For better land management, further field investigation to determine which scenario or combinations are substantial for the emergence of pathotrophs would be necessary.

Aboveground plant heterogeneity is linked to soil microbial taxonomic heterogeneity (Prober *et al*. 2015). In turn, maintaining microbial taxonomic heterogeneity may be achievable by cultivating various crops within and across sites. Several potential management strategies are available to promote aboveground plant diversity in agricultural systems (Funabashi 2018). However, the broader functional homogenization in our study underscores the importance of considering functionality alongside taxonomy when aiming for sustainable agriculture (Figure 3e, f). Future studies could incorporate chemical assessments of nutrient cycling functions, such as EcoPlate analysis or stable isotope techniques, to gain deeper insights into practical functionality (Miki *et al*. 2014; Oshiki *et al*. 2022).

## Conclusions

We compared soil microbial community structures and their predicted functions between natural lands and farmlands at within-site (∼200 m) to across-site (∼1500 km) scales in Kenya and Malawi. Our study revealed that farming activities drove the homogenization of soil microbial functional compositions more broadly than their taxonomic compositions, indicating consistent functional responses to agriculture even across locations with dissimilar taxa. Additionally, pathogenic fungi relatively increased in farmlands compared to natural lands, possibly due to decreased species competition and environmental conditions in the farmlands. While advancements in sequencing techniques have expanded our understanding of microbial taxonomic diversity, they may still overlook critical functional shifts. Our study highlights that incorporating microbial functional diversity is essential for accurately assessing the impacts of land-use changes on soil health.

## Endnotes

### Data and code availability

The sequence data used in this study is available in NCBI SRA PRJNA699079 and PRJNA699085, for soils collected in Kenya and Malawi, respectively. All scripts used in the analysis are available on GitHub (https://github.com/taka-ohi/KenyaMalawi_microbiome).

## Acknowledgements

We thank the local staff at the University of Nairobi and Lilongwe University of Agriculture and Natural Resources for their invaluable assistance with soil sampling and physical property measurements. We are also grateful to the Madegwa family for their generous support during the survey in Kenya, and to Mikiko Goto for her technical assistance with the molecular experiments. We also appreciate Enago (Crimson Interactive Japan Co., Ltd., www.enago.jp) for their professional English editing services. This research was supported by JSPS KAKENHI (grant numbers 18KK0183 and 21H02324) and Hokkaido University–Hitachi Cooperative Education and Research Support Program.

## Author contributions

YU and TO conceived the idea and designed the research. All authors contributed to sample collection. GK and KN conducted soil physical property analyses. TO performed chemical and molecular experiments, conducted DNA sequencing, analyzed the data, and wrote the first draft. All authors approved the final manuscript.

