## Supplementary materials for "Agricultural land use induces broader homogenization of soil microbial functional composition than taxonomic composition"

### Contents:

- **Figure S1**. Microbial taxa significantly changed the abundances by land use
- **Figure S2**. Dissimilarity of soil microbial functions in natural lands and farmlands
- **Figure S3**. Relationships between heterogeneities of environmental factors and microbial communities
- **Figure S4**. Contributions of fungal ASVs to community heterogeneity
- **Figure S5**. Composition of pathotrophs
- **Figure S6**. Relationship between soil moisture and $\beta$NTI
- **Figure S7**. Effect of DNA extraction buffer on microbial community structure
- **Table S1**. Pearson’s correlation between soil pH and relative abundances of pathogenic genera
- **Table S2**. Locations of the sampling sites
- **Table S3**. Soil texture
- **Table S4**. Buffer used for the DNA extraction and dilution rate for PCR/qPCR
- **Table S5**. KO number of functions related to Carbon and Nitrogen cycles used in this study

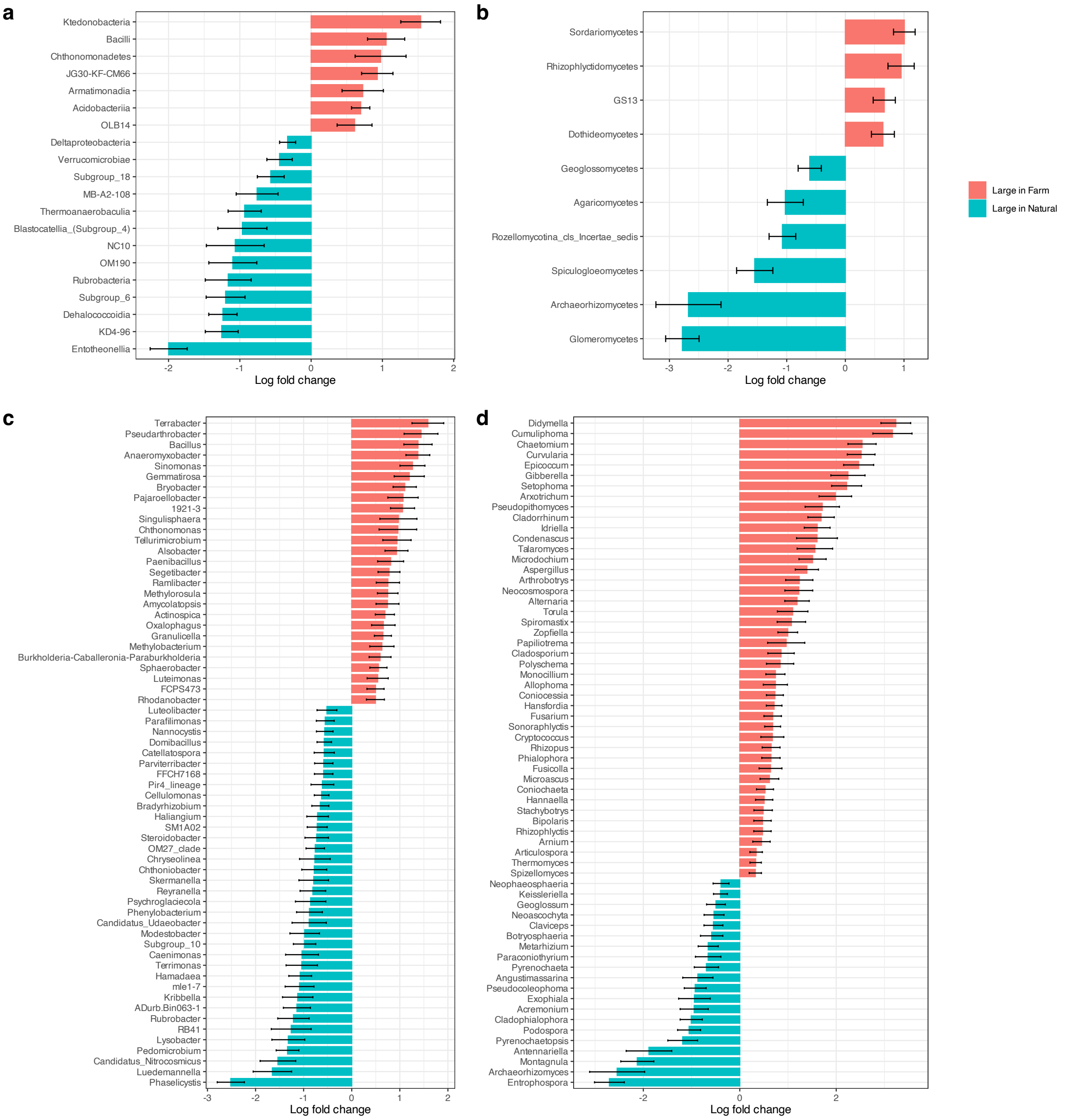


**Figure S1**. Microbial taxa significantly changed the abundances by land use. Taxa significantly changed their abundances between the land uses, tested by ANCOMBC were shown at class level (**a, b**) and genus level (**c, d**), for prokaryotes (**a, c**) and fungi (**b, d**). $Log_{10}$ fold changes of relative abundances in farmlands compared to natural lands across all sites are calculated. Microbial taxa that showed adjusted p-values < 0.05 are illustrated.

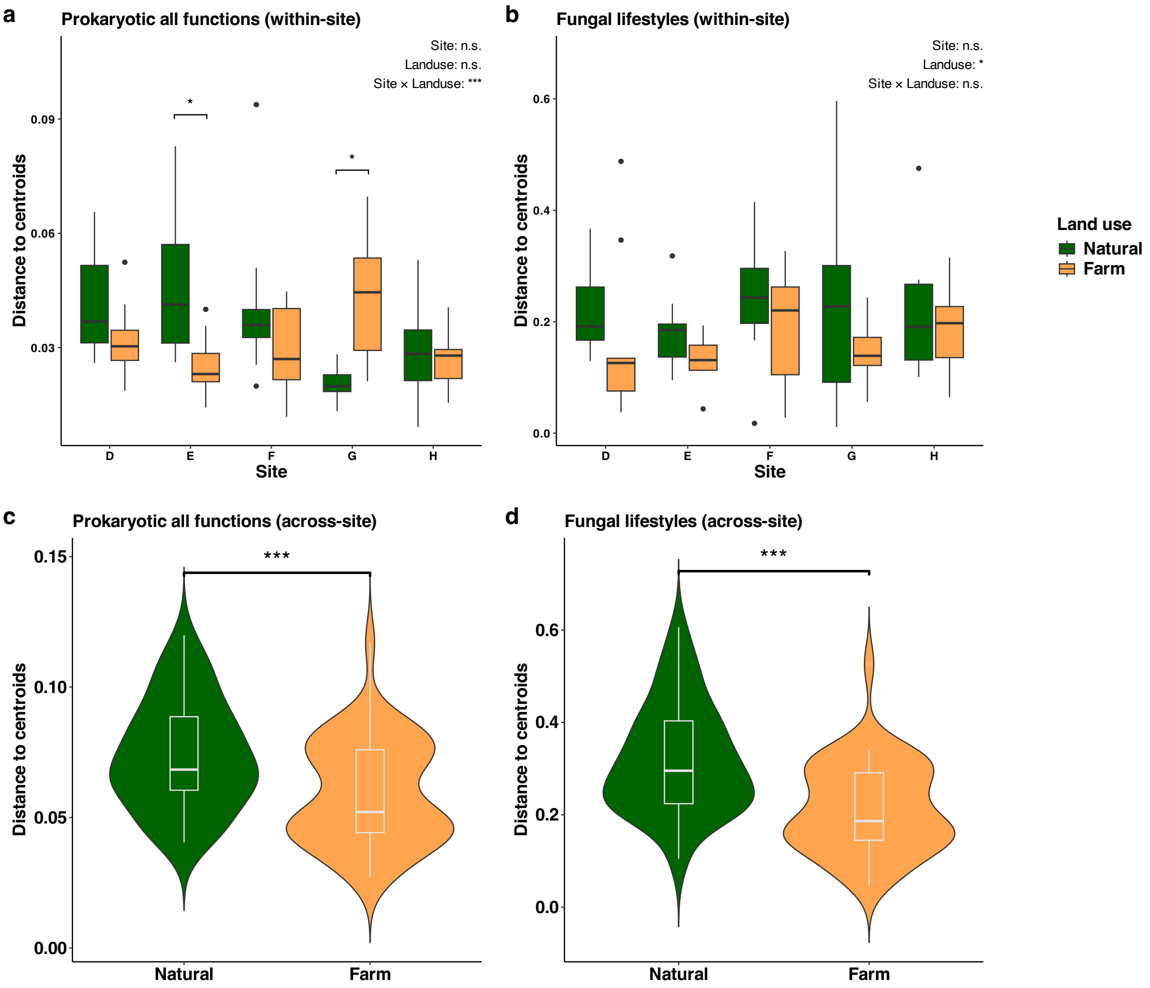


**Figure S2**. Dissimilarity of soil microbial functions in natural lands and farmlands. The distance to centroids of the functional compositions in each land use in (**c, d**) within-site and (**e, f**) across-site scales are plotted, for prokaryotes (**c, e**) and fungi (**d, f**). The p-values in the two-way ANOVA on sites, land uses and their interaction, and those in the t-test on land uses are indicated with "*", "**", or "***", representing $p < 0.05$, $p < 0.01$, or $p < 0.001$, respectively. If a significant interaction was found in the two-way ANOVA, pairwise comparisons of estimated marginal means were conducted to assess whether there were significant differences in land use within each site. Significant differences in land use within sites were indicated by a single asterisk "*".

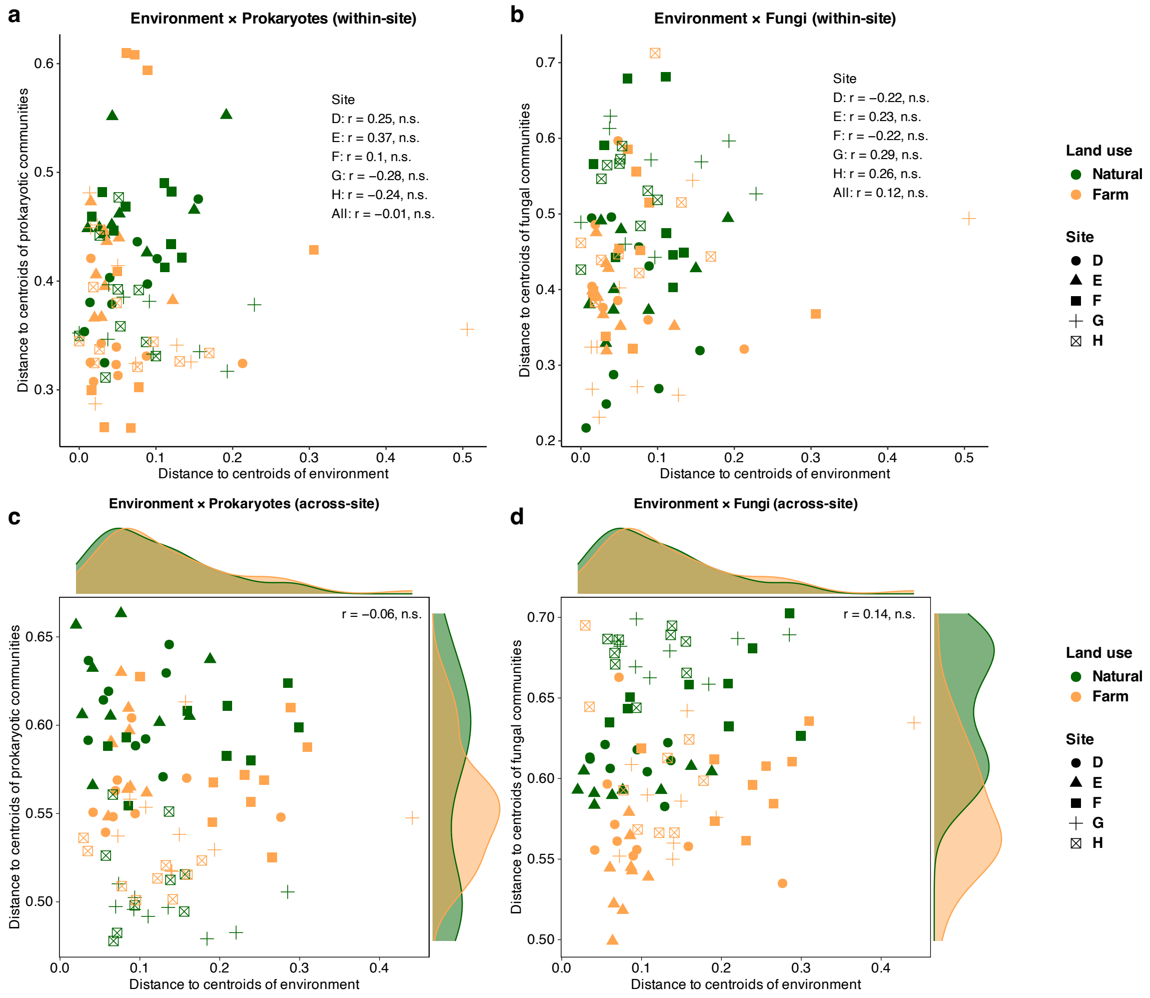


**Figure S3**. Relationships between heterogeneities of environmental factors and microbial communities. The correlations between distance to centroids of the environmental factors and that of microbial communities in each land use within site for (**a**) prokaryotes and (**b**) fungi, and across sites for (**c**) prokaryotes and (**d**) fungi are shown. The correlation coefficients and p-values in the Pearson's correlation tests are indicated.

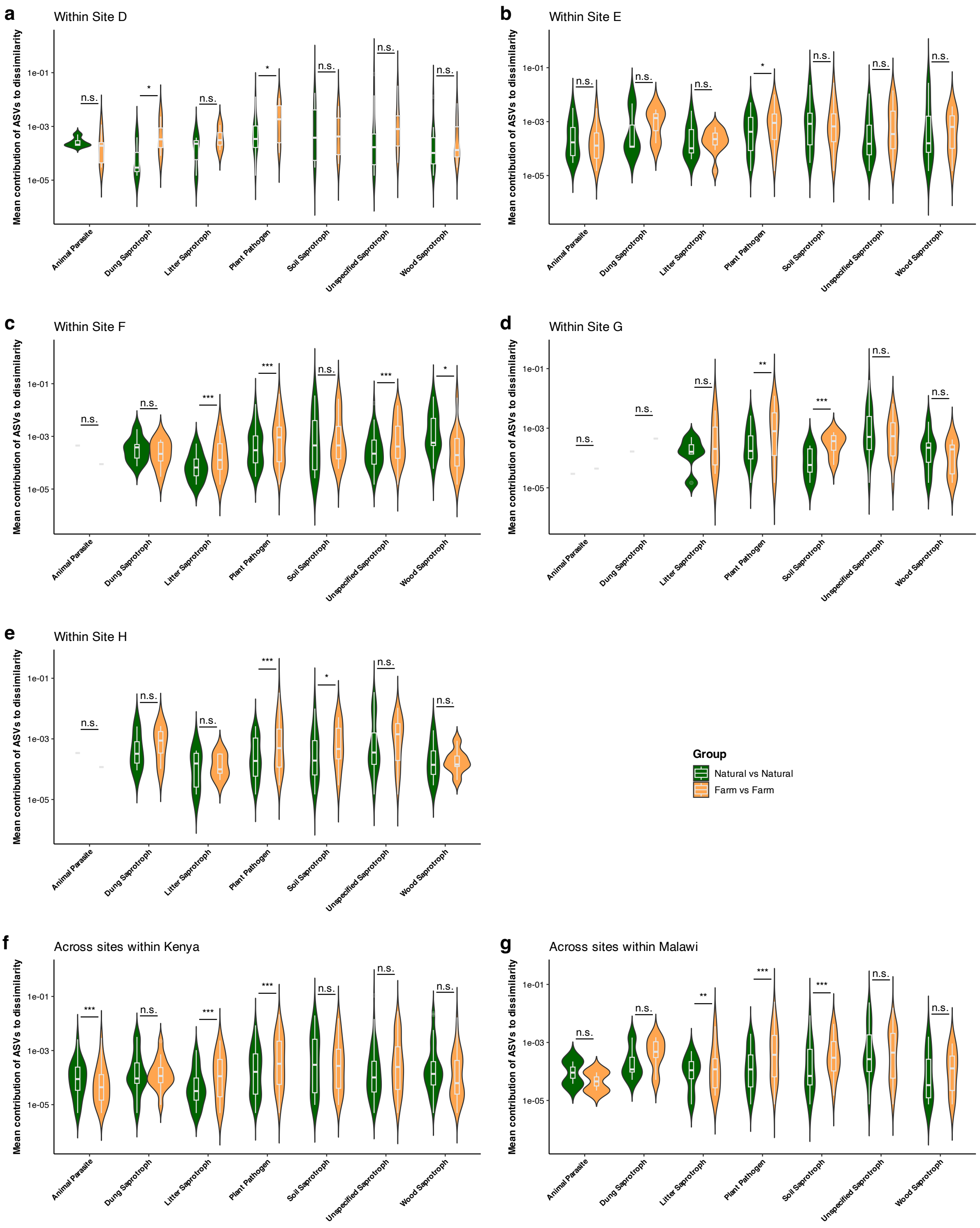


**Figure S4**. Contributions of fungal ASVs to community heterogeneity. The contributions of each fungal ASV to the Bray–Curtis dissimilarity among samples within each land use within site D–H (**a–e**) and within Kenya (**f**) and within Malawi (**g**) were averaged and grouped by fungal lifestyle. Asterisks ("*", "**", or "***") and n.s. indicate the significance levels of adjusted p-values of $< 0.05$, $p < 0.01$, $p < 0.001$, or no significant difference respectively, tested by 1000 permutations to assess differences in the mean values between natural lands and farmlands.

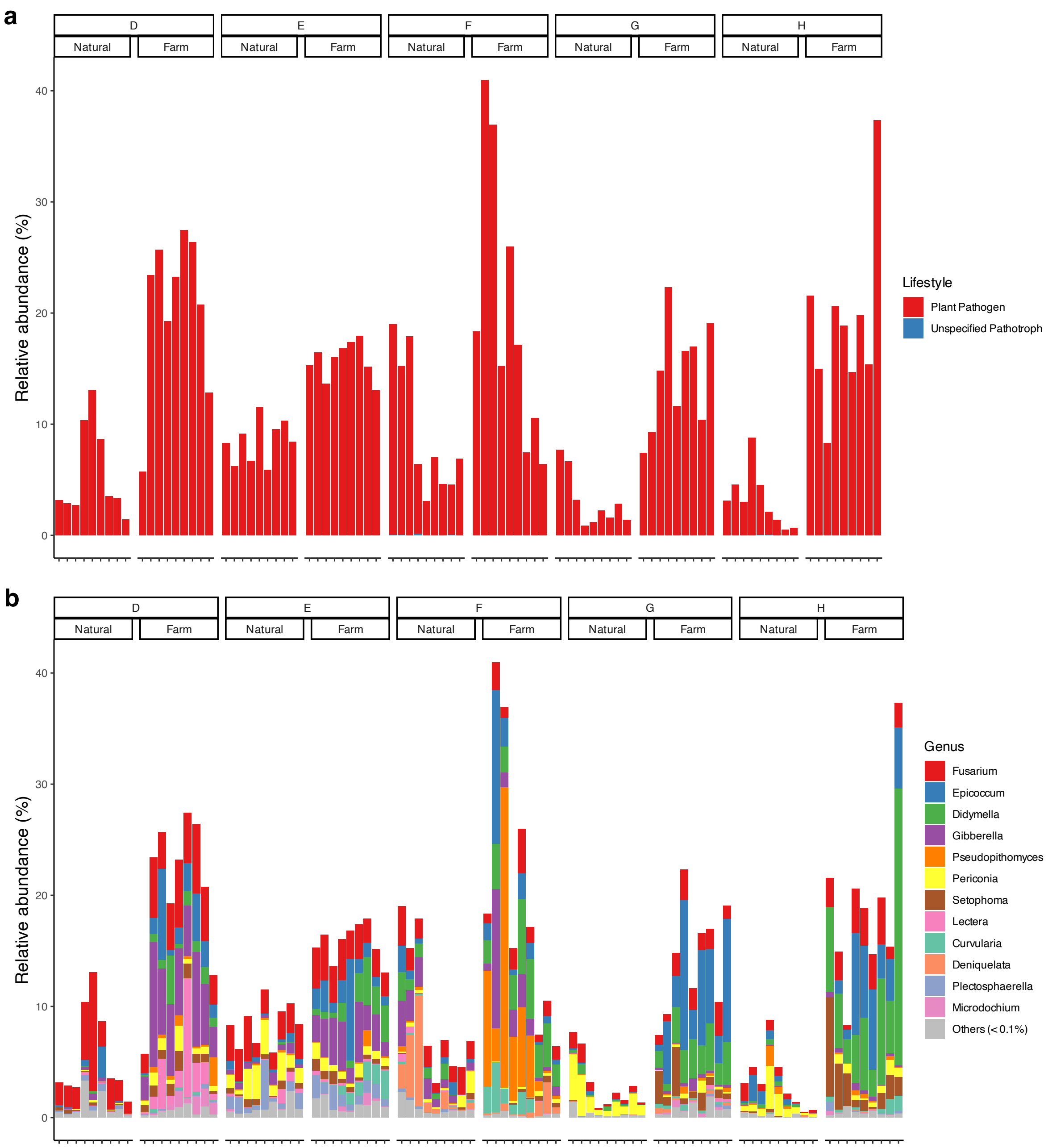


**Figure S5**. Composition of pathotrophs. The relative abundances of pathotrophs by primary lifestyles (**a**) and by genera that harbored mean abundances of > 0.1% (**b**) are shown.

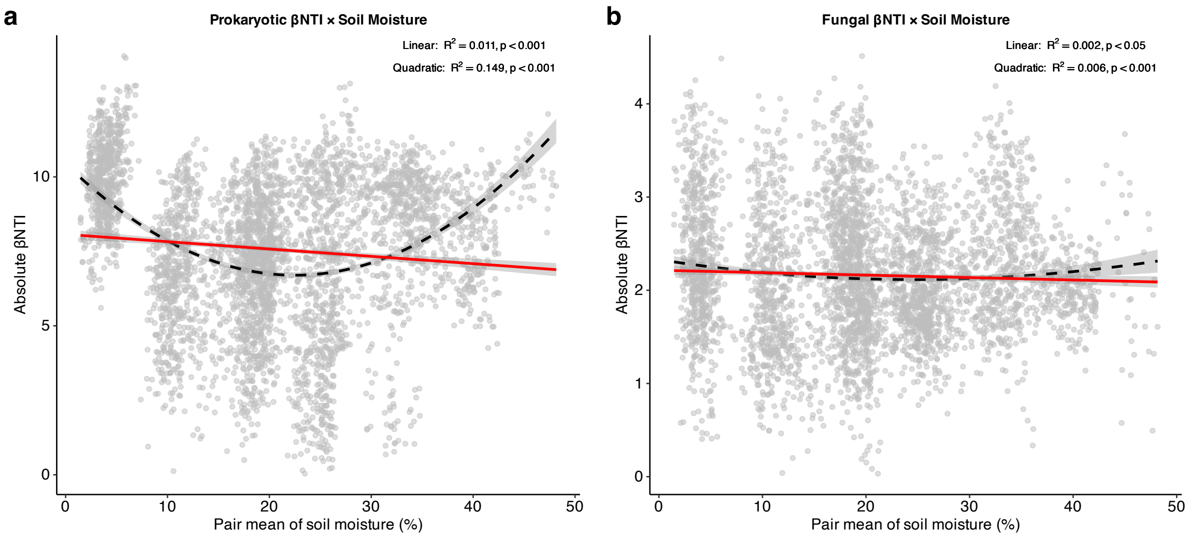


**Figure S6**. Relationship between soil moisture and $\beta$NTI. (**a**) Prokaryotic absolute $\beta$NTI and (**b**) fungal absolute $\beta$NTI are shown in relation to the mean soil moisture of corresponding sample pairs. The red solid line represents the linear regression model, while the black dashed line indicates the quadratic regression model. The gray shaded area corresponds to the 95% confidence interval. $R^2$ values and p-values for both linear and quadratic regressions are indicated.

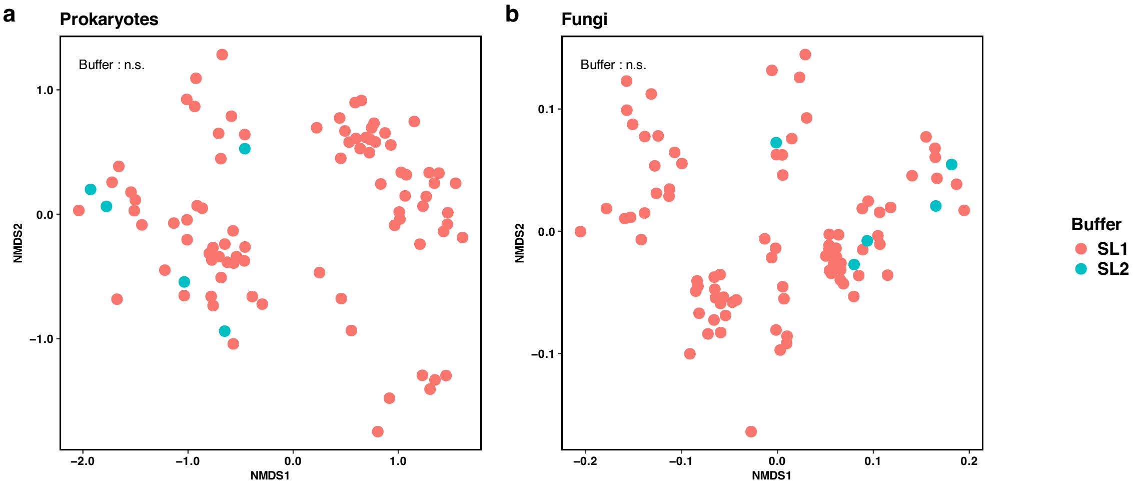


**Figure S7**. Effect of DNA extraction buffer on microbial community composition. NMDS for (**a**) prokaryotes and (**b**) fungi are shown. The p-values in the PERMANOVA are indicated with "n.s.", representing no significant difference.

**Table S1. Pearson's correlation between soil pH and relative abundances of pathotrophic genera**

| | **r** | **p** |
|---|---|---|
| *Fusarium* | –0.1 | n.s. |
| *Epicoccum* | –0.46 | < 0.001 |
| *Didymella* | –0.49 | < 0.001 |
| *Gibberella* | –0.18 | n.s. |
| *Pseudopithomyces* | –0.41 | < 0.001 |
| *Periconia* | 0.32 | < 0.01 |
| *Setophoma* | –0.31 | < 0.01 |
| *Lectera* | –0.03 | n.s. |
| *Curvularia* | –0.35 | < 0.001 |
| *Deniquelata* | 0.05 | n.s. |
| *Plectosphaerella* | 0.22 | < 0.05 |
| *Microdochium* | –0.09 | n.s. |
| *All* | –0.56 | < 0.001 |

**Table S2. Locations of the sampling sites**

| *Country* | *Site* | *Land use* | *Plot* | *Altitude* | *Latitude/Longitude* |
|---|---|---|---|---|---|
| Kenya | D | Farm | 1 | 1848 m | S1°14'50.59" E36°44'28.20" |
| Kenya | D | Farm | 2 | 1847 m | S1°14'50.03" E36°44'27.82" |
| Kenya | D | Farm | 3 | 1848 m | S1°14'49.52" E36°44'27.16" |
| Kenya | D | Natural | 1 | 1845 m | S1°14'47.21" E36°44'30.36" |
| Kenya | D | Natural | 2 | 1843 m | S1°14'47.12" E36°44'29.67" |
| Kenya | D | Natural | 3 | 1843 m | S1°14'47.04" E36°44'28.81" |
| Kenya | E | Farm | 1 | 1782 m | S1°15'28.18" E36°46'27.27" |
| Kenya | E | Farm | 2 | 1782 m | S1°15'28.28" E36°46'27.68" |
| Kenya | E | Farm | 3 | 1781 m | S1°15'28.23" E36°46'28.13" |
| Kenya | E | Natural | 1 | 1777 m | S1°15'27.40" E36°46'30.27" |
| Kenya | E | Natural | 2 | 1777 m | S1°15'27.75" E36°46'30.28" |
| Kenya | E | Natural | 3 | 1778 m | S1°15'28.07" E36°46'30.34" |
| Kenya | F | Farm | 1 | 1605 m | S1°34'54.45" E37°14'31.66" |
| Kenya | F | Farm | 2 | 1608 m | S1°34'54.21" E37°14'31.97" |
| Kenya | F | Farm | 3 | 1609 m | S1°34'54.15" E37°14'32.63" |
| Kenya | F | Natural | 1 | 1603 m | S1°34'52.16" E37°14'30.04" |
| Kenya | F | Natural | 2 | 1605 m | S1°34'52.00" E37°14'29.67" |
| Kenya | F | Natural | 3 | 1601 m | S1°34'51.30" E37°14'29.58" |
| Malawi | G | Farm | 1 | 1182 m | S14°11'10.78" E33°46'26.68" |
| Malawi | G | Farm | 2 | 1185 m | S14°11'10.73" E33°46'26.67" |
| Malawi | G | Farm | 3 | 1183 m | S14°11'14.98" E33°46'25.90" |
| Malawi | G | Natural | 1 | 1185 m | S14°11'15.78" E33°46'24.29" |
| Malawi | G | Natural | 2 | 1184 m | S14°11'14.85" E33°46'24.12" |
| Malawi | G | Natural | 3 | 1184 m | S14°11'15.09" E33°46'22.08" |
| Malawi | H | Farm | 1 | 1141 m | S13°58'24.74" E33°39'15.37" |
| Malawi | H | Farm | 2 | 1139 m | S13°58'26.10" E33°39'16.96" |
| Malawi | H | Farm | 3 | 1141 m | S13°58'23.71" E33°39'14.52" |
| Malawi | H | Natural | 1 | 1139 m | S13°58'25.88" E33°39'12.62" |
| Malawi | H | Natural | 2 | 1139 m | S13°58'25.55" E33°39'13.58" |
| Malawi | H | Natural | 3 | 1137 m | S13°58'25.62" E33°39'14.94" |

**Table S3. Soil texture**

| *Site* | *Land use* | *Sand (%)* | *Silt (%)* | *Clay (%)* |
|---|---|---|---|---|
| D | Natural | 38.67 ± 6.11 | 32.00 ± 2.00 | 29.33 ± 4.16 |
| D | Farm | 42.00 ± 2.00 | 32.67 ± 3.06 | 25.33 ± 3.06 |
| E | Natural | 39.33 ± 4.16 | 34.00 ± 2.00 | 26.67 ± 2.31 |
| E | Farm | 46.00 ± 8.72 | 26.67 ± 5.77 | 27.33 ± 3.06 |
| F | Natural | 53.33 ± 10.26 | 12.00 ± 9.17 | 34.67 ± 1.15 |
| F | Farm | 52.00 ± 5.29 | 14.00 ± 5.29 | 34.00 ± 2.00 |
| G | Natural | 68.33 ± 2.31 | 8.67 ± 2.31 | 23.00 ± 0.00 |
| G | Farm | 64.33 ± 2.31 | 11.00 ± 2.65 | 24.33 ± 0.58 |
| H | Natural | 64.33 ± 2.31 | 12.33 ± 0.58 | 24.33 ± 0.58 |
| H | Farm | 62.67 ± 0.58 | 12.33 ± 0.58 | 25.00 ± 0.00 |

**Table S4. Buffer used for the DNA extraction and dilution rate for PCR/qPCR**

| Sample | Site | Land use | Buffer | Dilution rate |
|---|---|---|---|---|
| K01 | D | Natural | SL1 | 500 |
| K02 | D | Natural | SL1 | 500 |
| K03 | D | Natural | SL1 | 500 |
| K04 | D | Natural | SL1 | 1 |
| K05 | D | Natural | SL2 | 100 |
| K06 | D | Natural | SL1 | 500 |
| K07 | D | Natural | SL1 | 500 |
| K08 | D | Natural | SL2 | 500 |
| K09 | D | Natural | SL1 | 500 |
| K10 | D | Farm | SL1 | 500 |
| K11 | D | Farm | SL1 | 100 |
| K12 | D | Farm | SL1 | 100 |
| K13 | D | Farm | SL1 | 1 |
| K14 | D | Farm | SL1 | 1 |
| K15 | D | Farm | SL1 | 50 |
| K16 | D | Farm | SL1 | 100 |
| K17 | D | Farm | SL2 | 100 |
| K18 | D | Farm | SL1 | 1 |
| K19 | E | Natural | SL1 | 100 |
| K20 | E | Natural | SL1 | 500 |
| K21 | E | Natural | SL1 | 500 |
| K22 | E | Natural | SL1 | 1 |
| K23 | E | Natural | SL1 | 50 |
| K24 | E | Natural | SL1 | 500 |
| K25 | E | Natural | SL1 | 10 |
| K26 | E | Natural | SL1 | 500 |
| K27 | E | Natural | SL2 | 1 |
| K28 | E | Farm | SL1 | 100 |
| K29 | E | Farm | SL1 | 1 |
| K30 | E | Farm | SL1 | 500 |
| K31 | E | Farm | SL1 | 100 |
| K32 | E | Farm | SL1 | 100 |
| K33 | E | Farm | SL1 | 1 |
| K34 | E | Farm | SL1 | 100 |

| | | | | |
|---|---|---|---|---|
| K35 | E | Farm | SL1 | 1 |
| K36 | E | Farm | SL1 | 1 |
| K37 | F | Natural | SL1 | 500 |
| K38 | F | Natural | SL1 | 500 |
| K39 | F | Natural | SL1 | 500 |
| K40 | F | Natural | SL1 | 50 |
| K41 | F | Natural | SL2 | 50 |
| K42 | F | Natural | SL1 | 500 |
| K43 | F | Natural | SL1 | 500 |
| K44 | F | Natural | SL1 | 500 |
| K45 | F | Natural | SL1 | 500 |
| K46 | F | Farm | SL1 | 10 |
| K47 | F | Farm | SL1 | 50 |
| K48 | F | Farm | SL1 | 50 |
| K49 | F | Farm | SL1 | 50 |
| K50 | F | Farm | SL1 | 50 |
| K51 | F | Farm | SL1 | 50 |
| K52 | F | Farm | SL1 | 50 |
| K53 | F | Farm | SL1 | 1 |
| K54 | F | Farm | SL1 | 50 |
| M01 | G | Natural | SL1 | 100 |
| M02 | G | Natural | SL1 | 100 |
| M03 | G | Natural | SL1 | 100 |
| M04 | G | Natural | SL1 | 100 |
| M05 | G | Natural | SL1 | 100 |
| M06 | G | Natural | SL1 | 100 |
| M07 | G | Natural | SL1 | 100 |
| M08 | G | Natural | SL1 | 100 |
| M09 | G | Natural | SL1 | 100 |
| M10 | G | Farm | SL1 | 1 |
| M11 | G | Farm | SL1 | 1 |
| M12 | G | Farm | SL1 | 10 |
| M13 | G | Farm | SL1 | 10 |
| M14 | G | Farm | SL1 | 1 |
| M15 | G | Farm | SL1 | 50 |
| M16 | G | Farm | SL1 | 50 |

| | | | | |
|---|---|---|---|---|
| M17 | G | Farm | SL1 | 50 |
| M18 | G | Farm | SL1 | 50 |
| M19 | H | Natural | SL1 | 100 |
| M20 | H | Natural | SL1 | 50 |
| M21 | H | Natural | SL1 | 100 |
| M22 | H | Natural | SL1 | 100 |
| M23 | H | Natural | SL1 | 100 |
| M24 | H | Natural | SL1 | 100 |
| M25 | H | Natural | SL1 | 10 |
| M26 | H | Natural | SL1 | 50 |
| M27 | H | Natural | SL1 | 50 |
| M28 | H | Farm | SL1 | 50 |
| M29 | H | Farm | SL1 | 10 |
| M30 | H | Farm | SL1 | 50 |
| M31 | H | Farm | SL1 | 50 |
| M32 | H | Farm | SL1 | 50 |
| M33 | H | Farm | SL1 | 10 |
| M34 | H | Farm | SL1 | 50 |
| M35 | H | Farm | SL1 | 100 |
| M36 | H | Farm | SL1 | 50 |

**Table S5. KO number of functions related to Carbon and Nitrogen cycles used in this study**

| KO | Symbol | Name | Cycle | Function | Pathway |
|---|---|---|---|---|---|
| K01601 | *rbcL, cbbL* | ribulose-bisphosphate carboxylase large chain [EC:4.1.1.39] | Carbon | Autotrophic C fixation | Calvin Benson |
| K01602 | *rbcS, cbbS* | ribulose-bisphosphate carboxylase small chain [EC:4.1.1.39] | Carbon | Autotrophic C fixation | Calvin Benson |
| K00239 | *sdhA, frdA* | succinate dehydrogenase flavoprotein subunit [EC:1.3.99.1] | Carbon | Autotrophic C fixation | rTCA |
| K00240 | *sdhB, frdB* | succinate dehydrogenase iron-sulfur subunit [EC:1.3.5.1] | Carbon | Autotrophic C fixation | rTCA |
| K00174 | *korA, oorA, oforA* | 2-oxoglutarate/2-oxoacid ferredoxin oxidoreductase subunit alpha [EC:1.2.7.3 1.2.7.11] | Carbon | Autotrophic C fixation | rTCA |
| K00175 | *korB, oorB, oforB* | 2-oxoglutarate/2-oxoacid ferredoxin oxidoreductase subunit beta [EC:1.2.7.3 1.2.7.11] | Carbon | Autotrophic C fixation | rTCA |
| K00176 | *korD, oorD* | 2-oxoglutarate ferredoxin oxidoreductase subunit delta [EC:1.2.7.3] | Carbon | Autotrophic C fixation | rTCA |
| K00177 | *korC, oorC* | 2-oxoglutarate ferredoxin oxidoreductase subunit gamma [EC:1.2.7.3] | Carbon | Autotrophic C fixation | rTCA |
| K15230 | *aclA* | ATP-citrate lyase alpha-subunit [EC:2.3.3.8] | Carbon | Autotrophic C fixation | rTCA |
| K15231 | *aclB* | ATP-citrate lyase beta-subunit [EC:2.3.3.8] | Carbon | Autotrophic C fixation | rTCA |
| K00192 | *cdhA* | anaerobic carbon-monoxide dehydrogenase, CODH/ACS complex subunit alpha [EC:1.2.7.4] | Carbon | Autotrophic C fixation | Wood-Ljungdahl |
| K00198 | *cooS, acsA* | anaerobic carbon-monoxide dehydrogenase catalytic subunit [EC:1.2.7.4] | Carbon | Autotrophic C fixation | Wood-Ljungdahl |
| K01964 | *NA* | acetyl-CoA/propionyl-CoA carboxylase [EC:6.4.1.2 6.4.1.3] | Carbon | Autotrophic C fixation | Hydroxypropionate Bicycle |

| | | | | | |
|---|---|---|---|---|---|
| K01965 | *PCCA, pccA* | propionyl-CoA carboxylase alpha chain [EC:6.4.1.3] | Carbon | Autotrophic C fixation | Hydroxypropionate Bicycle |
| K01966 | *PCCB, pccB* | propionyl-CoA carboxylase beta chain [EC:6.4.1.3 2.1.3.15] | Carbon | Autotrophic C fixation | Hydroxypropionate Bicycle |
| K11263 | *bccA, pccA* | acetyl-CoA/propionyl-CoA carboxylase, biotin carboxylase, biotin carboxyl carrier protein [EC:6.4.1.2 6.4.1.3 6.3.4.14] | Carbon | Autotrophic C fixation | Hydroxypropionate Bicycle |
| K01179 | *NA* | endoglucanase [EC:3.2.1.4] | Carbon | Cellulose breakdown | Cellulose breakdown |
| K01180 | *NA* | endo-1,3(4)-beta-glucanase [EC:3.2.1.6] | Carbon | Cellulose breakdown | Cellulose breakdown |
| K01225 | *CBH1* | cellulose 1,4-beta-cellobiosidase [EC:3.2.1.91] | Carbon | Cellulose breakdown | Cellulose breakdown |
| K01195 | *uidA, GUSB* | beta-glucuronidase [EC:3.2.1.31] | Carbon | Cellulose breakdown | Cellulose breakdown |
| K01181 | *xynA* | endo-1,4-beta-xylanase [EC:3.2.1.8] | Carbon | Xylan breakdown | Xylan breakdown |
| K01198 | *xynB* | xylan 1,4-beta-xylosidase [EC:3.2.1.37] | Carbon | Xylan breakdown | Xylan breakdown |
| K01183 | *NA* | chitinase [EC:3.2.1.14] | Carbon | Chitin breakdown | Chitin breakdown |
| K08689 | *bphAa, bphA1, bphA* | biphenyl 2,3-dioxygenase subunit alpha [EC:1.14.12.18] | Carbon | Lignin breakdown | Lignin breakdown |
| K15750 | *bphAb, bphA2, bph* | biphenyl 2,3-dioxygenase subunit beta [EC:1.14.12.18] | Carbon | Lignin breakdown | Lignin breakdown |
| K14582 | *nahB, doxE* | cis-1,2-dihydro-1,2-dihydroxynaphthalene/dibenzothiophene dihydrodiol dehydrogenase [EC:1.3.1.29 1.3.1.60] | Carbon | Lignin breakdown | Lignin breakdown |
| K03381 | *catA* | catechol 1,2-dioxygenase [EC:1.13.11.1] | Carbon | Lignin breakdown | Lignin breakdown |

| | | | | | |
|---|---|---|---|---|---|
| K14028 | *mdh1, mxaF* | methanol dehydrogenase (cytochrome c) subunit 1 [EC:1.1.2.7] | Carbon | Methane oxidation | Aerobic methane oxidation |
| K14029 | *mdh2, mxaI* | methanol dehydrogenase (cytochrome c) subunit 2 [EC:1.1.2.7] | Carbon | Methane oxidation | Aerobic methane oxidation |
| K00200 | *fwdA, fmdA* | formylmethanofuran dehydrogenase subunit A [EC:1.2.7.12] | Carbon | Methanogenesis | Methanogenesis |
| K00201 | *fwdB, fmdB* | formylmethanofuran dehydrogenase subunit B [EC:1.2.7.12] | Carbon | Methanogenesis | Methanogenesis |
| K00202 | *fwdC, fmdC* | formylmethanofuran dehydrogenase subunit C [EC:1.2.7.12] | Carbon | Methanogenesis | Methanogenesis |
| K00203 | *fwdD, fmdD* | formylmethanofuran dehydrogenase subunit D [EC:1.2.7.12] | Carbon | Methanogenesis | Methanogenesis |
| K11261 | *fwdE, fmdE* | formylmethanofuran dehydrogenase subunit E [EC:1.2.7.12] | Carbon | Methanogenesis | Methanogenesis |
| K00205 | *fwdF, fmdF* | 4Fe-4S ferredoxin | Carbon | Methanogenesis | Methanogenesis |
| K11260 | *fwdG* | 4Fe-4S ferredoxin | Carbon | Methanogenesis | Methanogenesis |
| K00204 | *fwdH* | 4Fe-4S ferredoxin | Carbon | Methanogenesis | Methanogenesis |
| K00672 | *ftr* | formylmethanofuran--tetrahydromethanopterin N-formyltransferase [EC:2.3.1.101] | Carbon | Methanogenesis | Methanogenesis |
| K01499 | *mch* | methenyltetrahydromethanopterin cyclohydrolase [EC:3.5.4.27] | Carbon | Methanogenesis | Methanogenesis |
| K00319 | *mtd* | methylenetetrahydromethanopterin dehydrogenase [EC:1.5.98.1] | Carbon | Methanogenesis | Methanogenesis |
| K00320 | *mer* | 5,10-methylenetetrahydromethanopterin reductase [EC:1.5.98.2] | Carbon | Methanogenesis | Methanogenesis |

| | | | | | |
|---|---|---|---|---|---|
| K00577 | *mtrA* | tetrahydromethanopterin S-methyltransferase subunit A [EC:2.1.1.86] | Carbon | Methanogenesis | Methanogenesis |
| K00578 | *mtrB* | tetrahydromethanopterin S-methyltransferase subunit B [EC:2.1.1.86] | Carbon | Methanogenesis | Methanogenesis |
| K00579 | *mtrC* | tetrahydromethanopterin S-methyltransferase subunit C [EC:2.1.1.86] | Carbon | Methanogenesis | Methanogenesis |
| K00580 | *mtrD* | tetrahydromethanopterin S-methyltransferase subunit D [EC:2.1.1.86] | Carbon | Methanogenesis | Methanogenesis |
| K00581 | *mtrE* | tetrahydromethanopterin S-methyltransferase subunit E [EC:2.1.1.86] | Carbon | Methanogenesis | Methanogenesis |
| K00582 | *mtrF* | tetrahydromethanopterin S-methyltransferase subunit F [EC:2.1.1.86] | Carbon | Methanogenesis | Methanogenesis |
| K00583 | *mtrG* | tetrahydromethanopterin S-methyltransferase subunit G [EC:2.1.1.86] | Carbon | Methanogenesis | Methanogenesis |
| K00584 | *mtrH* | tetrahydromethanopterin S-methyltransferase subunit H [EC:2.1.1.86] | Carbon | Methanogenesis | Methanogenesis |
| K00399 | *mcrA* | methyl-coenzyme M reductase alpha subunit [EC:2.8.4.1] | Carbon | Methanogenesis | Methanogenesis |
| K00401 | *mcrB* | methyl-coenzyme M reductase beta subunit [EC:2.8.4.1] | Carbon | Methanogenesis | Methanogenesis |
| K00402 | *mcrG* | methyl-coenzyme M reductase gamma subunit [EC:2.8.4.1] | Carbon | Methanogenesis | Methanogenesis |
| K01895 | *ACSS1_2, acs* | acetyl-CoA synthetase [EC:6.2.1.1] | Carbon | Methanogenesis | Methanogenesis |
| K00193 | *cdhC* | acetyl-CoA decarbonylase/synthase, CODH/ACS complex subunit beta [EC:2.3.1.169] | Carbon | Methanogenesis | Methanogenesis |

| | | | | | |
|---|---|---|---|---|---|
| K00197 | *cdhE, acsC* | acetyl-CoA decarbonylase/synthase, CODH/ACS complex subunit gamma [EC:2.1.1.245] | Carbon | Methanogenesis | Methanogenesis |
| K00194 | *cdhD, acsD* | acetyl-CoA decarbonylase/synthase, CODH/ACS complex subunit delta [EC:2.1.1.245] | Carbon | Methanogenesis | Methanogenesis |
| K14080 | *mtaA* | [methyl-Co(III) methanol/glycine betaine-specific corrinoid protein]:coenzyme M methyltransferase [EC:2.1.1.246?2.1.1.377] | Carbon | Methanogenesis | Methanogenesis |
| K04480 | *mtaB* | methanol---5-hydroxybenzimidazolylcobamide Co-methyltransferase [EC:2.1.1.90] | Carbon | Methanogenesis | Methanogenesis |
| K14081 | *mtaC* | methanol corrinoid protein | Carbon | Methanogenesis | Methanogenesis |
| K14082 | *mtbA* | [methyl-Co(III) methylamine-specific corrinoid protein]:coenzyme M methyltransferase [EC:2.1.1.247] | Carbon | Methanogenesis | Methanogenesis |
| K16176 | *mtmB* | methylamine---corrinoid protein Co-methyltransferase [EC:2.1.1.248] | Carbon | Methanogenesis | Methanogenesis |
| K16177 | *mtmC* | monomethylamine corrinoid protein | Carbon | Methanogenesis | Methanogenesis |
| K16178 | *mtbB* | dimethylamine---corrinoid protein Co-methyltransferase [EC:2.1.1.249] | Carbon | Methanogenesis | Methanogenesis |
| K16179 | *mtbC* | dimethylamine corrinoid protein | Carbon | Methanogenesis | Methanogenesis |
| K14083 | *mttB* | trimethylamine---corrinoid protein Co-methyltransferase [EC:2.1.1.250] | Carbon | Methanogenesis | Methanogenesis |
| K14084 | *mttC* | trimethylamine corrinoid protein | Carbon | Methanogenesis | Methanogenesis |
| K02588 | *nifH* | nitrogenase iron protein NifH | Nitrogen | N fixation | Nitrogen fixation |

| | | | | | |
|---|---|---|---|---|---|
| K02586 | *nifD* | nitrogenase molybdenum-iron protein alpha chain [EC:1.18.6.1] | Nitrogen | N fixation | Nitrogen fixation |
| K02591 | *nifK* | nitrogenase molybdenum-iron protein beta chain [EC:1.18.6.1] | Nitrogen | N fixation | Nitrogen fixation |
| K00531 | *anfG* | nitrogenase delta subunit [EC:1.18.6.1] | Nitrogen | N fixation | Nitrogen fixation |
| K22896 | *vnfD* | vanadium-dependent nitrogenase alpha chain [EC:1.18.6.2] | Nitrogen | N fixation | Nitrogen fixation |
| K22897 | *vnfK* | vanadium-dependent nitrogenase beta chain [EC:1.18.6.2] | Nitrogen | N fixation | Nitrogen fixation |
| K22898 | *vnfG* | vanadium nitrogenase delta subunit [EC:1.18.6.2] | Nitrogen | N fixation | Nitrogen fixation |
| K22899 | *vnfH* | vanadium nitrogenase iron protein | Nitrogen | N fixation | Nitrogen fixation |
| K10944 | *pmoA-amoA* | methane/ammonia monooxygenase subunit A [EC:1.14.18.3 1.14.99.39] | Nitrogen | Nitrification | Ammonia oxidation |
| K10945 | *pmoB-amoB* | methane/ammonia monooxygenase subunit B | Nitrogen | Nitrification | Ammonia oxidation |
| K10946 | *pmoC-amoC* | methane/ammonia monooxygenase subunit C | Nitrogen | Nitrification | Ammonia oxidation |
| K10535 | *hao* | hydroxylamine dehydrogenase [EC:1.7.2.6] | Nitrogen | Nitrification | Hydroxylamine oxidation |
| K00368 | *nirK* | nitrite reductase (NO-forming) [EC:1.7.2.1] | Nitrogen | Denitrification | Denitrification |
| K15864 | *nirS* | nitrite reductase (NO-forming) / hydroxylamine reductase [EC:1.7.2.1 1.7.99.1] | Nitrogen | Denitrification | Denitrification |
| K04561 | *norB* | nitric oxide reductase subunit B [EC:1.7.2.5] | Nitrogen | Denitrification | Denitrification |
| K02305 | *norC* | nitric oxide reductase subunit C | Nitrogen | Denitrification | Denitrification |
| K00376 | *nosZ* | nitrous-oxide reductase [EC:1.7.2.4] | Nitrogen | Denitrification | Denitrification |
| K00370 | *narG, narZ, nxrA* | nitrate reductase / nitrite oxidoreductase, alpha subunit [EC:1.7.5.1 1.7.99.-] | Nitrogen | Denitrification | Dissimilatory nitrate reduction |

| | | | | | |
|---|---|---|---|---|---|
| K00371 | *narH, narY, nxrB* | nitrate reductase / nitrite oxidoreductase, beta subunit [EC:1.7.5.1 1.7.99.-] | Nitrogen | Denitrification | Dissimilatory nitrate reduction |
| K00374 | *narI, narV* | nitrate reductase gamma subunit [EC:1.7.5.1 1.7.99.-] | Nitrogen | Denitrification | Dissimilatory nitrate reduction |
| K02567 | *napA* | nitrate reductase (cytochrome) [EC:1.9.6.1] | Nitrogen | Denitrification | Dissimilatory nitrate reduction |
| K02568 | *napB* | nitrate reductase (cytochrome), electron transfer subunit | Nitrogen | Denitrification | Dissimilatory nitrate reduction |
| K00362 | *nirB* | nitrite reductase (NADH) large subunit [EC:1.7.1.15] | Nitrogen | Denitrification | Dissimilatory nitrate reduction |
| K00363 | *nirD* | nitrite reductase (NADH) small subunit [EC:1.7.1.15] | Nitrogen | Denitrification | Dissimilatory nitrate reduction |
| K03385 | *nrfA* | nitrite reductase (cytochrome c-552) [EC:1.7.2.2] | Nitrogen | Denitrification | Dissimilatory nitrate reduction |
| K15876 | *nrfH* | cytochrome c nitrite reductase small subunit | Nitrogen | Denitrification | Dissimilatory nitrate reduction |
